# Endogenous intronic RNA tightly controls Cas9/CRISPR-mediated gene editing in human cells

**DOI:** 10.64898/2026.03.24.714022

**Authors:** André Lemos Carneiro, João Proença, Ehsan Valiollahi, Vasco M. Barreto

**Author notes:** To whom correspondence should be addressed. Vasco M. Barreto, André Lemos Carneiro.

## Abstract

CRISPR/Cas approaches are often limited by off-target effects. In vivo, multiple cell types are often present and off-targets may result from unintended targeting of the wrong cells. In this work, we propose a solution to this limitation by using a transcribed intron of the target gene as an endogenous trigger (intron triggers) for a novel conditional guide RNA (intcgRNA). In vitro, intcgRNAs were responsive to the presence of the trigger. As a proof-of concept, the human IL2 receptor subunit gamma gene (IL2RG) was then targeted using both the intcgRNA and the corresponding conventional crRNA in two cell lines: the lymphocytic HPB-ALL cell line, where IL2RG is highly expressed, and the epithelial HeLa cell line, where it is not. Sanger sequencing revealed that the crRNA and intcgRNA Cas9 complexes edited IL2RG with similar efficiency in HPB-ALL, whereas only the crRNA edited IL2RG in HeLa. This shows that intcgRNA avoids targeting unwanted cells that do not express the target gene, which is particularly relevant for in vivo targeting. The triggers of choice for conditional guides have been microRNAs, but as short intronic RNAs are far more diverse, trigger introns could become biomarkers of cell identity that improve the precision of CRISPR-based manipulations in vivo.

**GRAPHICAL ABSTRACT:** 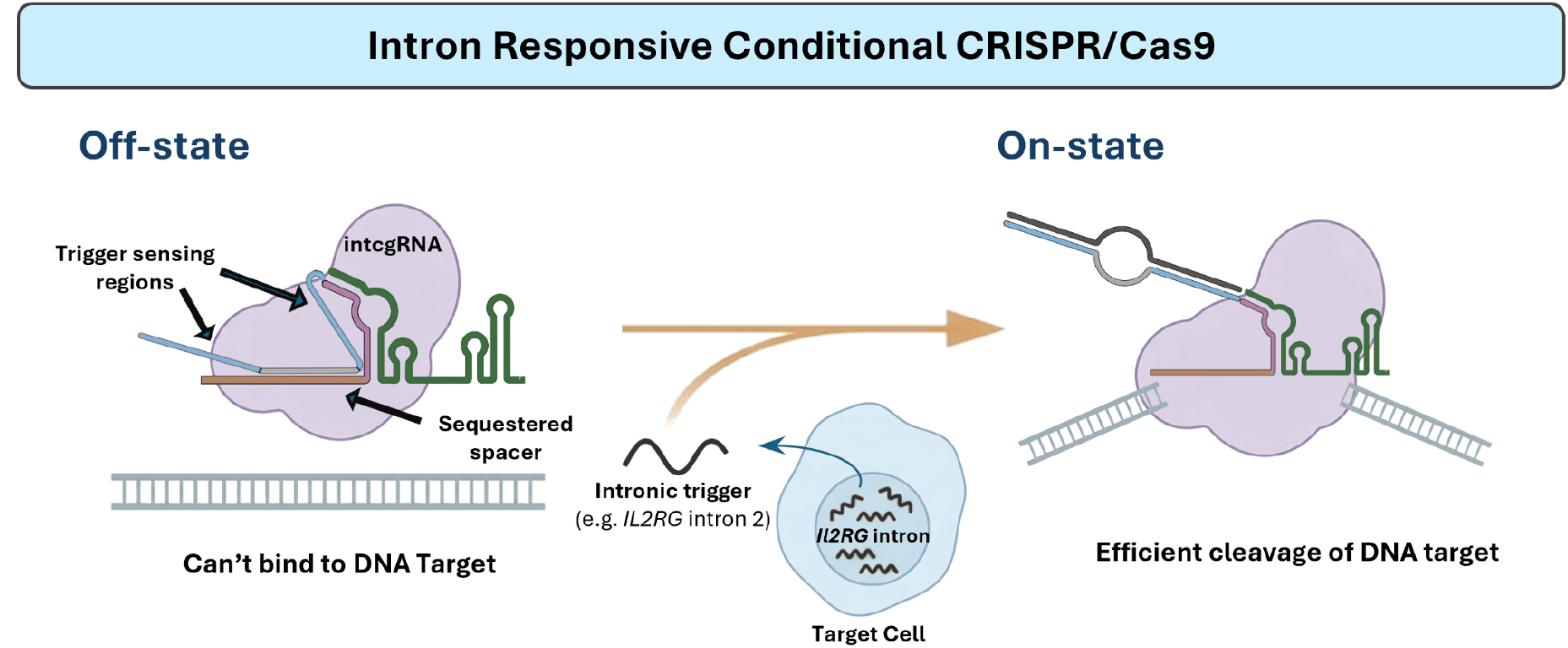

## INTRODUCTION

Gene editing was revolutionised in 2012-2013 with the demonstration that the components of a bacterial immune system against foreign nucleic acids, named Clustered regularly interspaced short palindromic repeats (CRISPR), cut DNA in vitro and the genome in human cells at precise locations (1–4). These components are a ribonucleoprotein (RNP) comprising an endonuclease (e.g., Cas9) and CRISPR RNAs, namely crRNA and trans-activating CRISPR RNA, which in artificial systems are often replaced by a single synthetic guide (sgRNA). The key component of the crRNA/sgRNA is the spacer region, which binds to a complementary sequence (the protospacer) in the genome, allowing the endonuclease to be targeted to the genomic region of choice.

Despite the enthusiastic adoption of the CRISPR technology for gene editing, which has reached clinical applications (e.g., (5)), problems remain, namely with respect to the delivery of the editing tool in vivo and the editing of unwanted regions in the genome (the off-targets). Unwanted sequences can be targeted because the protospacer is relatively small (20 nucleotides in CRISPR/Cas9) and tolerates mismatches, which creates many potential off-targets in the large vertebrate genomes. Restricting the levels of the RNP and the time window for editing reduces the off-targets (6). Furthermore, several ingenious approaches have been proposed, including the use of bioinformatic tools to find the guides with the fewest off-targets (7), the combined action of two Ca9-nickase-mutant RNPs that are guided by different sgRNAs to protospacers nearby in the genome, where they cooperate to produce the DSB (8), engineered Cas9 proteins with improved specificity (9–13), and methods that bypass the generation of DSBs, namely, the editing with base editors and prime-editing (14). These efforts have not completely solved the off-target problem. For instance, the improvements in specificity provided by the “high-fidelity” Cas9 variants are relatively small and specificity is negatively correlated with efficiency (15, 16). Likewise, base editors have been shown to have limited use (A-G and C-T) and to induce unwanted off-target mutations (17, 18).

Besides approaches to reduce off-targets by engineering the enzyme, the crRNA has also been manipulated. Chemical modifications, such as the incorporation of 2′-O-methyl-3′-phosphonoacetate at specific sites in the ribose-phosphate backbone, can significantly increase the specificity and flexibility of editing (19). Shorter spacers (with 17 to 19 nucleotides) have also been demonstrated to reduce off-target cleavage because they become more sensitive to mismatches (20) and even smaller spacers, which fail to induce editing, can be co-delivered with a normal 20-nucleotide spacer sgRNA to shield the off-targets (21). Furthermore, blocking the spacer sequence with a duplex and a hairpin structure was shown to interfere with cgRNA-DNA base pairing at off-target sites (22). More recently, as a subset of CRISPR/Cas conditional systems, crRNAs have been manipulated to render the RNP responsive to specific cellular signals or external stimuli (these crRNAs are known as conditional guide RNAs or cgRNAs). Examples include guides with RNA aptamers (aptazymes) (23), photocleavable groups in the spacer region (24) and the topologies that respond to the presence of RNA trigger molecules (25), allowing the cgRNA to switch state upon binding to its target ligand or trigger molecule.

Despite the growing interest in strategies employing cgRNAs, most still suffer from leakage and low performance in the active state compared to the native CRISPR/Cas system, and their implementation requires extensive optimisation, as the design rules are not yet fully understood. (26–28). Furthermore, for RNA trigger molecules, there is an upper size limit that imposes restrictions on the types of endogenous trigger molecules that can be used, which tend to be circumscribed to microRNAs.

Here, we show that a small intronic RNA from the target gene works as a trigger molecule to activate a newly designed cgRNA, which we named intcgRNA. The data show that Cas9 is active only in cells that strongly express the target gene, which is relevant for in vivo targeting strategies. The use of intronic RNA as a trigger could vastly increase the potential number of biomarker trigger molecules that circumscribe CRISPR activity to the target cell type in vivo.

## MATERIAL AND METHODS

### In Vitro Cas9 Cleavage Assay

To evaluate the cleavage activity for the native and conditional gRNAs. First, 11 μL of mixture A was prepared, containing 500 nM of crRNA or intcgRNA (either one annealed to tracrRNA using the procedure described below), 250-2000 nM of trigger RNA molecule, 1 μL of RiboLock RNase Inhibitor (Thermo Fisher), and Duplex Buffer (Integrated DNA Technologies). Mixture A was incubated at 37ºC for 60 min. Next, 15 μL of a reaction mixture B containing 1X NEBuffer 3.1 (NEB), 500 nM Cas9 (Integrated DNA Technologies), 1.5 μL of linearized px330-mSA plasmid(1.7 μg, Addgene plasmid #113096 (29)) with the spacer target sequence in RNase-Free Water (Nzytech) was added into mixture A and the resulting mixture was incubated at 37ºc for 4 hours. The reaction mixture was sampled at different time points. After collection, 8 μL of Q buffer (0.1 mg/mL Proteinase K, 0.01 M Tris-HCl pH 8.3, 0.000144% SDS) and 1 μL of 0.5 M EDTA (Panreac AppliChem) were added to each sample to digest DNA-bound Cas9 and inactivate Cas9, respectively. Each sample was also incubated at 65ºC to guarantee further reaction inactivation. The reaction mixture was then analysed using 1.3% agarose gel electrophoresis. For tests where RNPs were assembled before trigger sensing, native gRNA/intcgRNAs were incubated with Cas9 in duplex buffer with the RiboLock RNase Inhibitor for 20 minutes before the trigger molecule was added to the mixture.RNPs were then incubated with the trigger molecule for 1 hour prior to being added to the mixture with the 1X NEBuffer 3.1(NEB) and linearized target plasmid.

### Cloning of the IL2RG target sequence into px330-mSA and plasmid linearization

Oligos sequences: 2ACc.TargetIL2RGFw (5’ CACCTGGTAATGATGGCTTCAACATGG) and 2ACc.TargetIL2RGRv (5’AAACCCATGTTGAAGCCATCATTACCA) were annealed to insert the *IL2RG* region sequence (including PAM) targeted by the intcgRNA into px330-mSA. The plasmid px330-mSA was digested with the BbsI restriction enzyme (New England Biolabs) for 30 minutes at 37°C. The digested plasmid was then purified using a QIAquick Gel Extraction Kit (Qiagen) and eluted in EB buffer. The sgRNA oligos were phosphorylated and annealed by mixing 1 μL of each oligo (100 μM), 1 μL of 10X T4 DNA Ligase buffer (ABclonal, 2,000,000 U/ml), 6.5 μL of ddH2O, and 0.5 μL of T4 PNK (NEB) in a total volume of 10 μL. The oligo mixture was incubated at 37°C for 30 minutes, heated to 95°C for 5 minutes, and then ramped down to 25°C at 5°C/min using a thermocycler. The ligation reaction was set up by combining 50 ng of BbsI-digested pX330 plasmid, 1 μL of phosphorylated and annealed oligo duplex (1:200 dilution), 5 μL of T4 ligase buffer, 0.5 μL T4 DNA Ligase (ABclonal), and ddH2O to a total volume of 10 μL. T4 DNA Ligase was added to the mixture, and the reaction was incubated at room temperature for 4 hours. Finally, competent cells were transformed with the ligated plasmid, and positive clones were screened using colony PCR and sequencing. For use in the in vitro cleavage assay, the sequence-verified px330-mSA plasmid was linearized by digestion with EcoRV(NEB).

### Heat shock transformation of Ultra-Competent *E. coli*

Fifty μL of Ultra-Competent DH5α *E. coli* were added to 10 μL of the ligation reaction and kept on ice for 30 minutes. The mixture was then heat-shocked at 42°C for 1 minute and 30 seconds, followed by incubation on ice for 2 minutes. 800 μL of LB medium was added to the mixture, and the cells were incubated at 37°C for 1 hour. Following incubation, the transformed cells were plated onto selective antibiotic agar plates.

### intcgRNA design optimization

The trigger-sensing regions of the intcgRNA were optimised using NUPACK design tool. The input window for trigger choice was the intron 2 sequence of *IL2RG*. Sequence patterns: AAAA, CCCC, GGGG, UUUU, KKKKKK, MMMMMM, RRRRRR, SSSSSS, WWWWWW,YYYYYY, were prohibited. Candidate trigger sequences were additionally screened using blastn on the NCBI BLAST web server to minimise undesired sequence complementarity with the intcgRNA. intcgRNA/gRNA crRNAs and trigger sequences used are shown in Supplementary Table 1.

### Cell culture procedures

Cells were cultured at 37ºC, 5% CO2. HPB-all cells were grown in Roswell Park Memorial Institute medium (RPMI 1640, Biowest), supplemented with 10% heat-inactivated Foetal Bovine Serum (Biowest), and Penicillin/Streptomycin (Gibco) at 100 μ/mL. HeLa cells were grown in Minimum Essential Medium (Biowest), supplemented with 10% heat-inactivated Foetal Bovine Serum (Biowest) and Penicillin/Streptomycin (Gibco) at 100 μ/mL.

### Transfection

All cell lines were transfected by electroporation. Briefly, the 1-2 million cell pellet was resuspended in 100 μL of electroporation buffer (5 mM KCl, 15 mM MgCl2, 120 mM Na2HPO4/ NaH2PO4, pH 7.2, 50 mM Mannitol, and 0.05% PEG). 0.104 nmol of Cas9(IDT), 0.12 nmol of crRNA or intcgRNA crRNA (either one annealed to tracrRNA using the same procedure described below (without addition of RNase Inhibitor), were added to the reaction mix. The mix was transferred to an electroporation cuvette and electroporated in an Amaxa Nucleofector IIb using the program X-001 for all cell lines. Afterwards, 200 μL of culture medium was added to the cuvette and the cells transferred to cell culture plates. Cells were analysed by flow cytometry 72 hours after nucleofection.

### Annealing of RNA Oligos

The RNA oligos were obtained dry (lyophilized) and resuspended in IDTE buffer (IDT) to a concentration of 100 μM. For annealing, equal volumes of oligos were mixed and diluted in Duplex Buffer (IDT; 14 μL of Duplex Buffer for every 2 μL of native crRNA/tracrRNA or intcrRNA/tracrRNA mixture), then incubated at 95 °C for 4 minutes. The samples were then gradually cooled to 25 °C at a rate of 1 °C/min. If necessary, they were further diluted to the desired concentration in Duplex Buffer.

### Oligo Synthesis

All RNA and DNA single-strand oligos were purchased from Integrated DNA Technologies.

### DNA sequencing

STABVIDA performed Sanger sequencing.

### Flow Cytometry

To analyse cells by flow cytometry, they were first harvested and pelleted by centrifugation at 300 rcf for 5 minutes. The cell pellets were resuspended in PBS containing 1% FBS and washed twice. Cells were analyzed on a Sony SH800Z. GFP was measured with a 488 nm laser and a 525/50 band-pass filter. The propidium iodide (PI) signal was collected using a 488 nm laser and a 600/60 band-pass filter. At least 5000 events in live-cell/singlet gates were recorded for analysis. Genomic DNA of sorted cells was extracted immediately.

### Genomic DNA extraction

To purify genomic DNA, the cell pellet was resuspended in 200 μL of Q buffer (as previously described). The suspension was subjected to the following thermocycler program: 50ºC for 2 hours, 95ºC for 15 minutes.

### Reverse transcription quantitative real-time PCR (qRT-PCR)

Cells were centrifuged at 300×g for 5 min at room temperature, and total RNA was extracted from cell pellets using tripleXtractor (GRiSP) according to the manufacturer’s instructions. To remove residual genomic DNA, RNA was treated with DNase I (Ambion) according to the manufacturer’s protocol prior to reverse transcription. Total RNA was reverse transcribed using NZY M-MuLV Reverse Transcriptase (NZYTech) with random hexamers, following the manufacturer’s instructions; RiboLock RNase inhibitor (Thermo Fisher Scientific) was used in place of the NZY ribonuclease inhibitor. Reverse transcription reactions were performed in a Biometra Gradient thermocycler.

qPCR amplification was performed on a LightCycler 480 (Roche) real-time PCR system using SYBR GreenER qPCR SuperMix for ABI PRISM (Thermo Fisher Scientific). Primers were used at a final concentration of 100 nM each: *IL2RG* forward 5′-GTGGAAGTGCTCAGCATTGG-3′ and reverse 5′-ACAGAGATAACCACGGCTTCC-3′; β-actin (*ACTB*) forward 5′-CCAACTGGGACGACATGGAG-3′ and reverse 5′-GGGTGTTGAAGGTCTCAAAC-3′.. qPCR conditions included an initial denaturation at 95 °C for 10 min followed by 45 cycles of 95 °C for 15 s, 57 °C for 15 s, and 72 °C for 10 s, with fluorescence acquisition at 72 °C. Relative expression was calculated using the 2−ΔΔCt method, normalizing *IL2RG* to *ACTB*.

### Quantifying Indel%

Synthego ICE (30) was used to infer CRISPR edits from Sanger trace data. The primers used to sequence the target region were IL2RGFw (5’-AGTCAAGGAAGAGGCATGGC) and IL2RGRv (5’-AGATCTCTGTACGGCCCCTT).

### Statistical Analysis

All statistical analyses were performed using GraphPad Prism 10.6.1. For each experimental condition, cells were independently seeded and transfected in three independent biological replicates. Data are presented as mean ± standard deviation (SD). Editing efficiencies between cell lines and guide RNA conditions were compared using two-way ANOVA followed by Šidák’s multiple comparisons test. Significance levels are indicated as follows: two symbols, p < 0.01; three symbols, p < 0.001; four symbols, p < 0.0001.

### Genome-wide analysis of mRNA and intron length distributions

Genome-wide data for Figure 4 were obtained from Homo_sapiens.GRCh38.115.gtf (Ensembl release 115). mRNA length distribution was calculated for protein-coding genes using the smallest transcript per gene as the representative transcript. Intron length distribution was calculated using all annotated transcripts, after removal of duplicate introns to retain unique introns across isoforms. Short introns were defined as introns ≤300 nt. Genes containing at least one such intron were classified as short-intron genes. OMIM disease genes containing short introns were identified by direct Ensembl gene ID matching to the OMIM gene list. The number of annotated human miRNAs used for comparison was obtained from miRBase v22.

## RESULTS

### Design of intcgRNA, a novel RNA inducible cgRNA triggered by intronic RNA

Attempts at strand-displacement-based activation or deactivation of guides with RNA have used short messenger RNA or miRNA (25). We reasoned that intronic RNA could be an alternative source of trigger molecules because, as most genes have at least one short intron, the ability to target specific cell types based on their signature transcriptome would increase. Furthermore, whereas messenger RNA and miRNA are found dispersed throughout the nucleus (32, 33), introns tend to be spliced and degraded in the general proximity of the transcription site (34), potentially giving an intron a high spatial CRISPR-Cas9 control within the nucleus of the targeted cell. To test an intronic RNA trigger, we initially developed a novel allosterically activatable cgRNA based on Galizi et al.’s design (35). Two key design modifications were introduced (Figure 1). First, the extension was relocated from the 5’ end of the gRNA to the 3’ upper stem of the crRNA molecule. This change enabled greater flexibility in determining the blocking duplex’s location. This also circumvented the contradictory findings surrounding 5’ extensions: some studies report that a duplex placed outside the targeting sequence at the 5’ end suppresses gRNA activity upon trigger binding (27), while others using nearly identical designs find no such inhibition (35, 36). Given these inconsistencies, avoiding the 5’ end altogether offered a more reliable design strategy. Furthermore, an uncharacterized mechanism has been shown to cleave the 5’ extensions of gRNAs in human cells (8, 37), thereby increasing design complexity. Second, the duplex region was shifted from the PAM-distal end of the spacer to the PAM-proximal end. This modification aimed to enhance the performance of the off-to-on state transition by blocking the seed region, a critical step in gRNA cleavage activity. The design retained the duplex and sensing region sizes from Galizi et al.’s original design, i.e., 13 and 15 base pairs, respectively. Upon the trigger molecule binding to the trigger sensing region, a conformational change exposes the spacer without the trigger directly binding to it, as in cgRNAs that can be freely programmed to sense triggers that are unrelated to the spacer sequence (27).

**Figure 1.**
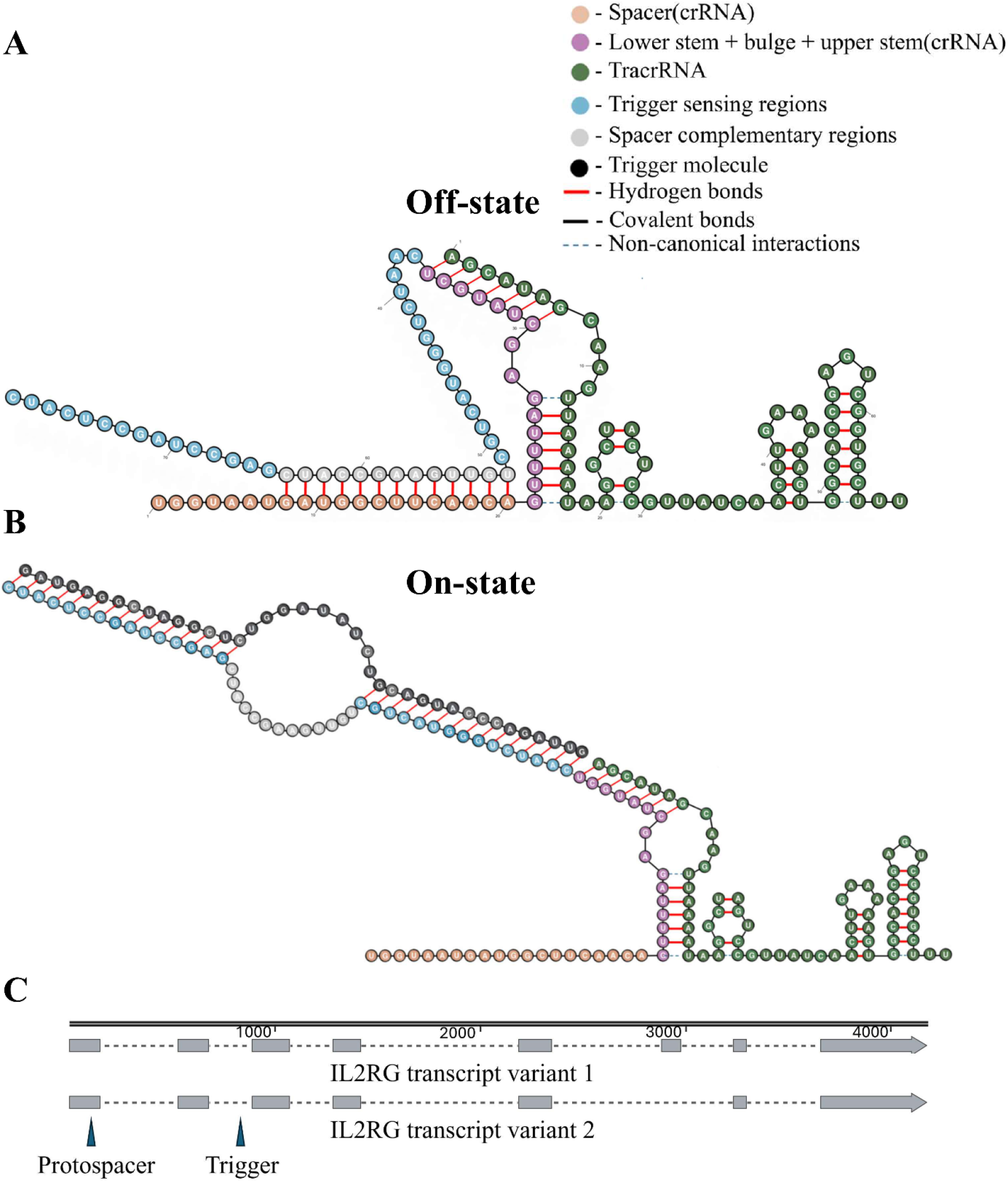
Novel allosteric cgRNA applied to a spacer targeting *IL2RG* (28), activatable by a sequence from an *IL2RG* intron with a size of 208 bp. The sub-sequence of the intron used for activation was selected with the aid of NUPACK (43). **A)** The novel intcgRNA in its off-state, where cleavage is expected to be abrogated. **B)** On-state, where intcgRNA is expected to bind to target and promote cleavage with activation done by an intronic trigger molecule that binds to the trigger sensing regions and exerts a conformational change that unwinds the duplex present in the spacer sequence based on Galizi et.al (35) **C)** Locations in the *IL2RG* locus of the target sequence and the intron encoding the trigger molecule.

For a proof of principle that intronic RNA sequences can be used as trigger molecules, we established criteria for selecting an appropriate gene target: 1) The gene must contain at least one small intron (< 300 bp) to ensure efficient cgRNA activation, as the activation performance decreases rapidly with trigger length; (36) 2) The gene should have high expression in at least one cell line and very low or no expression in another to serve as a trigger-absent control; the expression threshold is set at >110.4 normalised transcripts per million (nTPM) based on the successful case of cgRNA activation with the survivin gene in HeLa cells (36) and expression data from The Human Protein Atlas Project database (website) (38,39); 3) A high-efficiency gRNA for the target gene encoding the intron must have been reported in the literature. A Python scraper script was created to extract data from the Human Protein Atlas database. The interleukin-2 receptor gamma gene (*IL2RG*) was selected as the target. (41–43), meeting our criteria, as it has high expression in HPB ALL cells (526.1 nTPM) and negligible expression in HeLa cells (0.1 nTPM). The 208-bp intron 2 of *IL2RG* was chosen as the trigger molecule. The NUPACK web server was then used to identify a thermodynamically favourable 39-nucleotide sequence within this intron for cgRNA activation(Supplementary Figure were sorted 72 hours after electrop1(43).

### In vitro, intcgRNA is tightly controlled by the trigger

To assess if the newly proposed design(with a 13-bp duplex region) would work, an in vitro cleavage assay was first performed. Briefly, Cas9 was added to a solution containing the intcgRNA, the trigger RNA molecule, and a linear plasmid with the target sequence, and the final mixture was incubated at 37ºc and then analysed by agarose gel electrophoresis to detect the Cas9-induced cleavage of the plasmid (Figure 2A). The in vitro cleavage assay showed that the proposed design worked as intended in a cell-free environment. The cleavage product was not observed in the absence of the trigger RNA molecule. Remarkably, even after 7 hours of digestion, no cleavage was detected in the off state, thus validating the design’s functionality (data not shown). However, the trigger was left to incubate with the cgRNA before adding Cas9, which does not mimic the in vivo context, where the intcgRNA and Cas9 will be complexed before meeting the trigger RNA molecule. Furthermore, Hunt et al. 2022 noticed that the cgRNA they tested had difficulty sensing the trigger RNA molecule after it was complexed with Cas9 (28). Thus, the Cas9 in vitro cleavage assay was repeated but this time the cgRNA and Cas9 RNPs were formed prior to the incubation with the trigger RNA molecule (Figure 2B). Despite a significant hindrance of cleavage activity on the cgRNA complexed in RNPs before being incubated with the trigger, the cgRNA could still function. In an attempt to facilitate the cgRNA activation, we then experimented with diminishing the size of the duplex from 13 bp to 12 and 11 nucleotides and tested them using the in-vitro cleavage assay. As the duplex region length diminished, the cleavage activity increased because displacement becomes easier given the reduced number of base pairs in the duplex, resulting in weaker binding (Figure 2C). However, we could also observe that decreasing the duplex length to 12 and 11 base pairs, while improving cleavage activity, also caused the cgRNA to show cleavage activity in its off state, becoming leaky. The cgRNA with the smallest duplex had the most cleavage activity in the off state, which appeared similar to the one from the cgRNA with a 13 bp duplex in the on state. These data validate the overall design, but leave open the question whether the cgRNA will work efficiently in cells.

**Figure 2.**
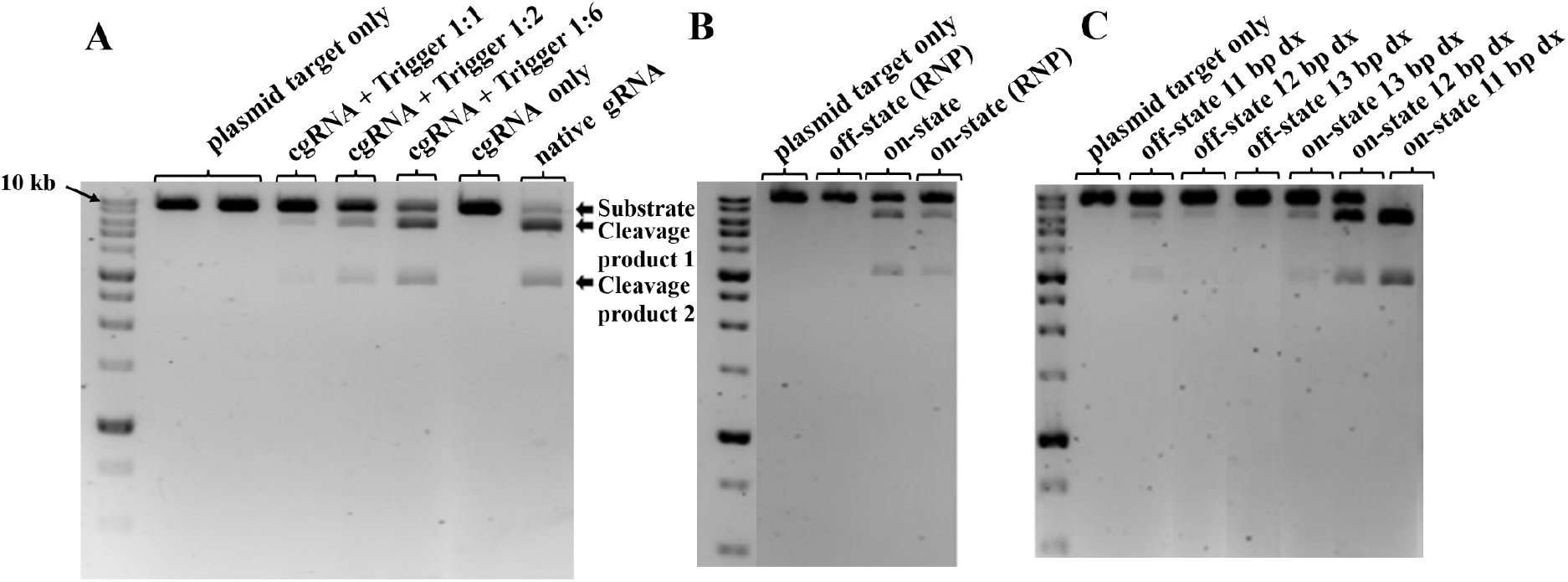
Cas9 in vitro cleavage assay to evaluate intcgRNA edits the target region in vitro with an efficiency similar to the conventional crRNA in the presence of the trigger. A) Cas9 cleavage after 127 minutes when the trigger was incubated with the intcgRNA with a 13 bp spacer duplex, prior to Cas9 being added to the mixture. The progressive increase in cleavage product formation over time and with increasing trigger RNA quantities (in the tested ratios range) demonstrates the correct functioning of the designed intcgRNA. B) Cas9 in vitro cleavage assay when the intcgRNA with a 13 bp spacer duplex and Cas9 form an RNP prior to the incubation with the trigger RNA molecule. C) Influence of spacer duplex length on off state(intcgRNA+Cas9) and on state (intcgRNA+Cas9 + trigger) performance when RNPs are formed prior to the addition of the trigger. Cas9 cleavage after 127 minutes. For all experiments, the samples were run on a 1.8% or 2.0% agarose gel alongside a 10 kb ladder (PCRBIO DNA Ladder II) for size comparison. Cas9 cleavage of the linearized plasmid substrate (9,046 bp) yields two expected fragments of approximately 6098 bp (cleavage product 1) and 2948 bp (cleavage product 2). The two products produced by Cas9 cleavage are marked with arrows.

### intcgRNA restricts editing activity to cells expressing the trigger

To test the performance of intcgRNAs in cells, we first confirmed by qRT-PCR that the published data on the expression of *IL2RG* are correct, as the expression of the gene in HPB-ALL is more than two orders of magnitude higher than in HeLa cells (Figure 3A). Three conditional gRNAs were then delivered via RNP electroporation to the HeLa and HPB-ALL human cell lines, and the transfected (GFP+) cells were sorted 72 hours after electroporation (Supplementary Figure 2, Supplementary Tables 2 and 3 for sorting and purity data). The editing efficiencies were measured on the purified cells based on Sanger sequencing data using Synthego ICE (30). As expected, the native crRNA produced the high levels of editing in the HPB-ALL and HeLa cell lines (Figure 3B and C). In contrast, whereas intcgRNA induced editing in the HPB-ALL cells, in HeLa, which does not express the trigger, the cgRNAs with 11-bp and 12-bp duplex in the spacer showed no detectable indel activity according to Synthego ICE analysis (Figure 3B and C). Furthermore, an analysis of the indels created by Cas9 showed that, in HPB-ALL, the intcgRNA is both quantitatively and qualitatively similar to the standard gRNA (Supplementary Figure 3). Interestingly, the results deviated from initial expectations based on the in vitro observations. The guide that showed no activity in the off state in the in vitro cleavage assay paradoxically consistently produced residual indels in cells(< 10%). Conversely, the cgRNAs that exhibited more leaky activity in vitro produced little to no detectable cleavage in cells. These results show that certain versions of the intcgRNA effectively restrict Cas9 to cells expressing the trigger RNA, achieving editing efficiencies above 60% in HPB-ALL while remaining undetectable by Sanger sequencing in HeLa..

**Figure 3.**
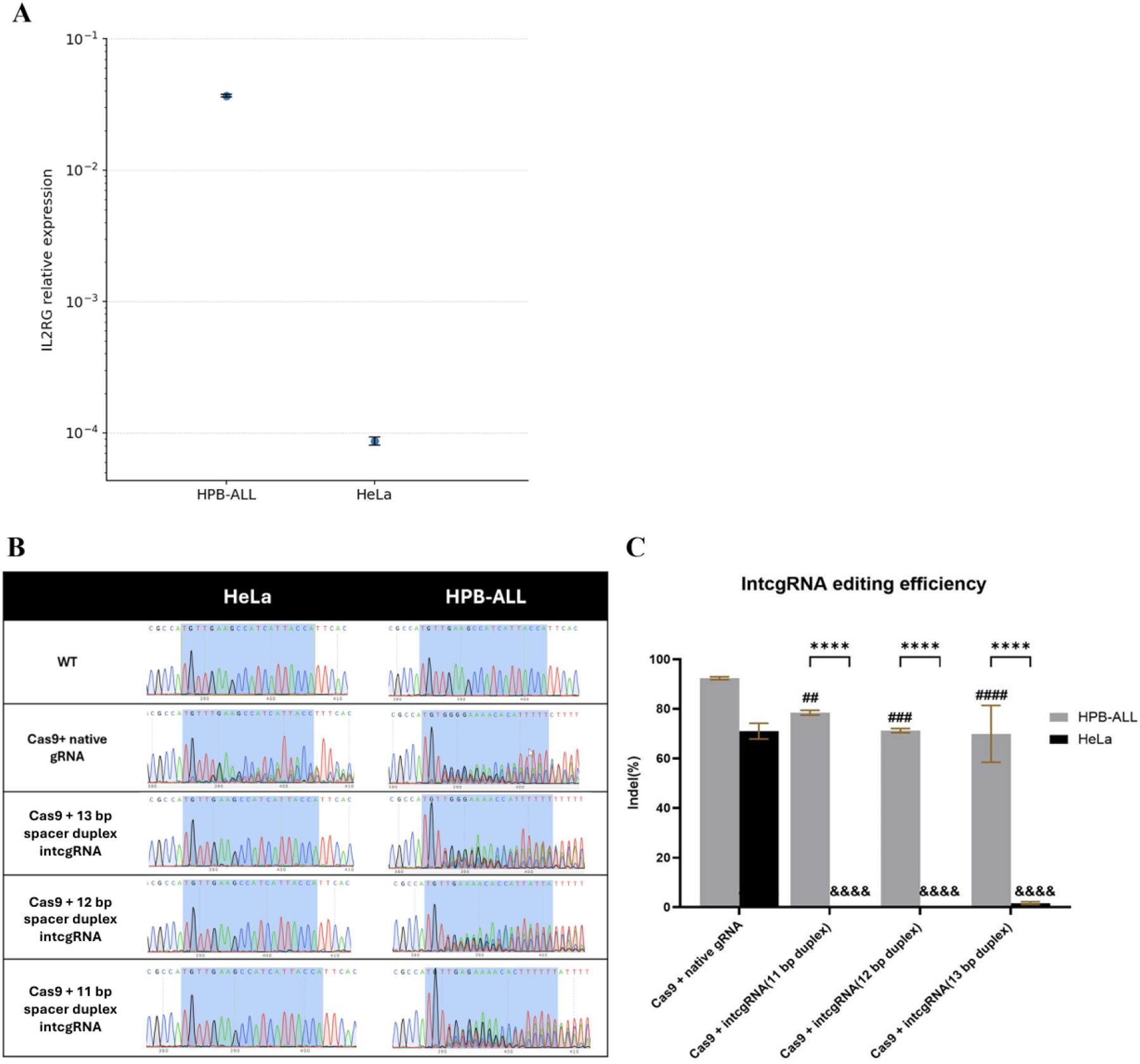
intcgRNA edits the target with an efficiency similar to that of the conventional crRNA, but only in cells in which the target gene is markedly transcribed. A) RNA expression levels of IL2RG transcripts in HeLa and HPB-ALL cells normalized to β-actin, detected by RT-qPCR. B) Sequence alignments of the target region amplified from GFP+ cells 72 hours after electroporation with the RNP complexes and a GFP-encoding plasmid. C) Quantification of the editing events detected by Sanger sequencing using ICE (30). The data from B and C are representative of two independent experiments. Bars show mean ± SD (n = 3 independent biological replicates). Statistical significance was determined using two-way ANOVA followed by Šídák’s multiple comparisons test. Symbols denote significance of comparisons: * indicates comparison between HPB-ALL and HeLa for the same experimental condition; # indicates comparison of each intcgRNA to native gRNA condition within HPB-ALL; & indicates comparison of each intcgRNA to native gRNA condition within HeLa. Symbol key: two symbols, p < 0.01; three symbols, p < 0.001; four symbols, p < 0.0001.

### Genome-wide evaluation of potential intronic RNA as trigger molecules

Despite the tightness of intcgRNA, this system will only be truly helpful if intron triggers expand the regulatory possibilities created by other classes of RNA triggers, namely microRNAs. It is known that mRNA data are better at discriminating cells than microRNA data (45, 46) and there is an obvious correlation between the number of mRNAs and that of introns. Furthermore, a genome-wide analysis based on *Homo_sapiens*.*GRCh38*.*115*.*gtf* identified 56,912 unique short introns (≤300 nt) among 415,845 unique introns across all annotated transcripts, and 12,843 genes with at least one short intron, representing 63.8% of the 20,121 protein-coding genes analysed. This greatly exceeds the number of human microRNAs (1,917 in miRBase v22; (47)). Thus, there is an emerging trend to explore the enormous potential of intronic RNAs as biomarkers of cell identity (48).In Figure 4, we compare the abundance and size distributions of intronic RNAs and microRNAs as potential CRISPR triggers. Panel A shows the length distribution of human mRNAs, providing context for intron sizes. Panel B displays the distribution of intron lengths, highlighting the abundant population of short introns (≤300 nt) that are functionally suitable for intcgRNA activation. Panel C compares the number of genes containing short introns (12,843 genes, representing 63.8% of protein-coding genes) with the total number of human microRNAs (1,917). Direct matching to the OMIM disease-gene dataset further showed that 11,292 of 17,408 OMIM genes (64.9%) contain at least one short intron that could serve as a trigger.

**Figure 4.**
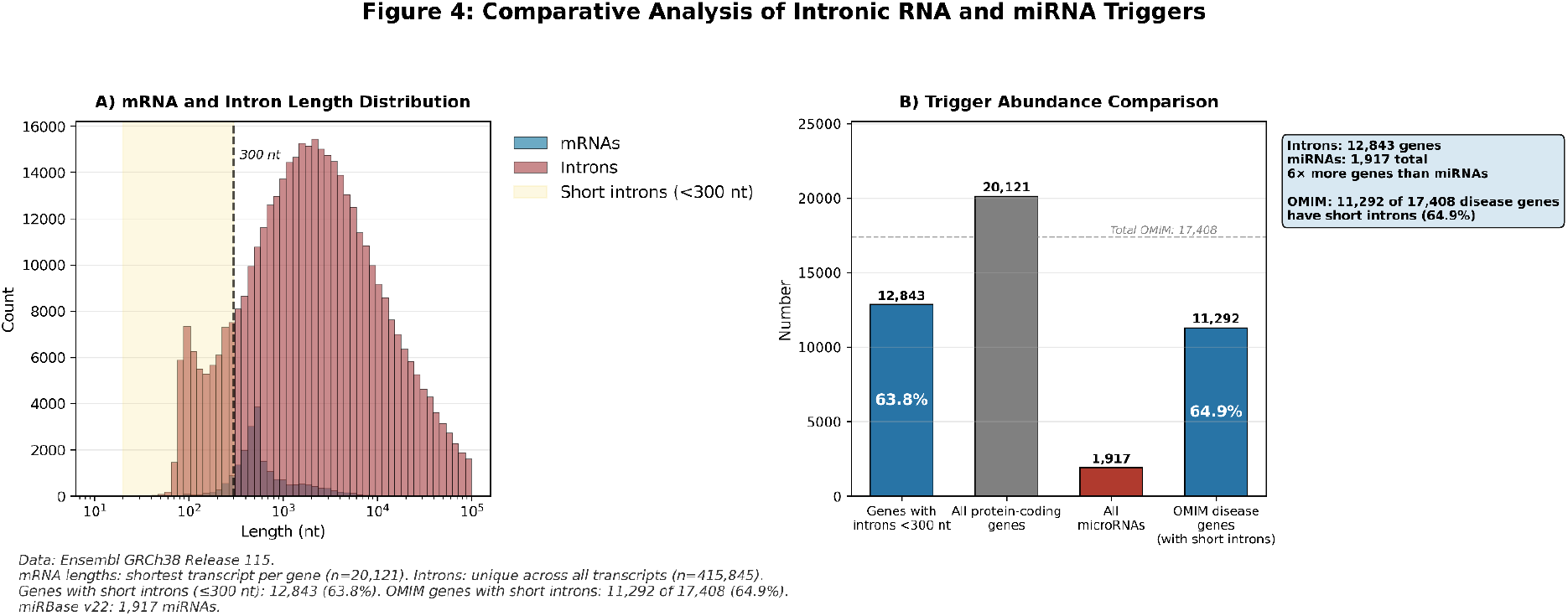
Short intronic RNAs are a diverse source of endogenous trigger molecules for CRISPR-based manipulations. **A)** Distribution of mRNA and unique intron lengths across all annotated transcripts lengths for human protein-coding genes, using the smallest transcript per gene as the representative transcript; the shaded region indicates short introns (≤300 nt). **B)** Comparison of trigger abundance, showing the number of protein-coding genes with at least one short intron (12,843), the total number of protein-coding genes (20,121), the total number of annotated human miRNAs (1,917), and the number of disease-associated genes with at least one short intron (11,292 of 17,408). Percentages indicate the fraction of all protein-coding genes or OMIM disease genes containing at least one short intron. Data were derived from *Homo_sapiens*.*GRCh38*.*115*.*gtf* (Ensembl release 115) and miRBase v22. For intron analyses, all transcripts were considered and duplicate introns were removed.

These data suggests that intronic RNA triggers have enormous potential as biomarkers that circumscribe gene editing to specific cell types and, potentially, limit editing activity to the target locus, if an intronic RNA from that locus activates the CRISPR-intcgRNA and the activation is local and does not reach off-targets.

## Discussion

Here we describe the first conditional CRISPR guides that respond to intronic RNA, establishing a new class of trigger molecules with the potential to produce cell-specific gene editing. This study contributes to a large body of work on conditional gRNAs, which were first reported by Tang et al. in 2017 (23). Since then, most reports on conditional gRNAs have focused on in vitro data (37, 49), data in bacteria (35, 50–52) or data in eukaryotic cells using artificial triggers (synthetic molecules or transcripts from transgenes) and/or artificial target genes (transgenes) (38, 53–57). However, the list of studies showing gene editing activity in endogenous targets (unmanipulated genomic sequences) from systems using endogenous RNA triggers is growing. (25, 58–63). Strikingly, in all studies from this last group, microRNA triggers were used, sometimes as alternatives to mRNA triggers. Hunt and Chen concluded that the full-length mCherry mRNA translocates into the cytosol too fast to function as an efficient trigger (28), whereas Wang et al. mention the structural diversity of mRNA as a handicap (61). To be assertive about the causes of the limitations of RNAs as triggers, one would need a systematic study evaluating how editing efficiency depends on the RNA class, length, and expression levels of the trigger molecule. However, the prevalence of microRNA triggers suggests that empirical observations from independent researchers have led to this choice. Because of their small size, many intronic RNAs are also an alternative to this problem, as they can be used in place of the corresponding mRNA, and are potentially a better choice than microRNAs because of their higher diversity.

In our study, the intronic RNA is derived from the target gene because pharmacological and genetic studies have established that intronic RNAs are spliced and cleared very close to the gene locus (34, 64–66). Thus, in addition to the cell-specific editing we show in this report, this design has the potential to confine editing activity to the target locus. However, many reports have challenged the locus-associated view by showing that several intron-derived RNAs are stable, abundant and not confined to the gene locus, including stable circular lariats, which can be found throughout the nucleus (67) or the cytoplasm (68), intronic RNAs that function as precursors of microRNAs (Mirtrons) (69) or stable intronic sequence RNAs (sisRNAs) in the nucleus (70), of which the linear Full-Length Excised Introns (FLEXIs) are a subset (48, 71). In the future, this diversity of behaviours could be leveraged. For instance, locus-associated intronic RNAs could be the primary choice if off-targets are indeed avoided and the target gene is expressed in the target cells, as in, for instance, the engineering of CAR-T and CAR-NK cells, the targeting of genes involved in lineage commitment in hematopoietic stem and progenitor cells, and gene therapy of autologous somatic cells and differentiated tissues. In contrast, stable intronic RNAs from genes other than the target one that are found throughout the nucleus or/and cytoplasm would be used as trigger molecules, just like microRNAs, if the main goal is cell-specific identity and the target cell does not express the target gene, as in most of the targeting of induced pluripotent cells, which typically involves editing a gene that is expressed after differentiation, or the editing of regulatory regions, which are not transcribed.

As a preliminary report on a new class of RNA triggers, this study has several limitations. First, we have not directly shown that the trigger is indeed the intron molecule and not the pre-mRNA, some shorter aborted transcript or a transcript retaining the intron. It is also possible that these species are not mutually exclusive and that each contributes to activation, as any RNA containing the trigger-complementary sequence could, in principle, bind the sensing region of the intcgRNA. However, several lines of evidence make these alternatives unlikely. The *IL2RG* pre-mRNA exceeds 4 kb, and trigger activation efficiency decreases sharply with RNA length (37), consistent with reports that full-length mRNAs are poor triggers due to size and structural complexity (28, 61). Prematurely terminated transcripts are rapidly degraded by the nuclear RNA exosome (72–74). As for intron retention, there is no evidence of *IL2RG* isoforms retaining intron 2 (IL2RG ENSG00000147168, (77)), and at 208 bp, this intron lies within the size range associated with intron definition, which favours efficient splicing (78). Even if a fraction of transcripts did retain it, nonsense-mediated decay would limit their steady-state abundance relative to the excised intron pool. Taken together, the simplest and most likely explanation is that the excised intron 2 of *IL2RG* is the primary functional trigger.

Second, we describe only one target gene, and this approach needs to be extended to more candidates to determine if it is practical. Third, although our new intcgRNA is extremely effective and tight, we have barely explored the space of possibilities. Notably, the results in vitro and in cells were not fully consistent, as the tightest guide in the in vitro cleavage assay was the one that produced more indels in the absence of the trigger. It is common for guides to show high cleavage efficiency in tube assays but poor cleavage in cells (69,70), but we have no simple explanation for why, in cells, the intcgRNA with the longer duplex region is the leakiest one, which is paradoxical. One possible explanation is that the longer trigger-sensing/blocking duplex region is more susceptible to partial hybridization by cellular RNAs. Although these RNAs may be unable to bind to the full sensing/blocking region and fully activate the intcgRNA, partial binding could still destabilize the blocking duplex sufficiently to permit the residual editing observed. Fourth, we have postponed the analysis of the off-targets and the use of next-generation sequencing to increase the resolution of the data, because these analyses are not essential for the proof-of-principle and require time and resources. Thus, given the complexity of intronic RNAs in terms of cellular location, half-life, and topology, and the open question raised by the unexplored topology of the intcgRNAs, only a larger systematic approach will identify the ideal features of intcgRNA and the criteria to identify the best candidates.

This study introduces intronic RNA as a new source of trigger molecules for CRISPR-based approaches. We used Cas9-based gene editing, but this new tool could be adopted by the sister technologies that use base editors or other CRISPR endonucleases, and beyond gene editing. Future studies will address whether the locus-associated intronic RNA restricts the activity of intcgRNA to the target locus, thus diminishing the indels in the off-targets and whether the plethora of FLEXIs and other forms of stable intronic RNAs can be leveraged for CRISPR-based cell-specific manipulations.

## ACKNOWLEDGEMENTS

The authors would like to thank Zoé Enderlin Vaz da Silva for her help with the cell-sorting experiments. We are also grateful to Bianca Miranda Basso, Diogo Magalhães, Miguel Chaves-Ferreira, Nadiya Kubasova, and Zoé Enderlin Vaz da Silva for sharing their expertise, experimental protocols, and troubleshooting advice.

## AUTHOR CONTRIBUTIONS

André Lemos Carneiro: Conceptualization, Investigation, Methodology, Data curation, Formal analysis, Validation, Writing—original draft.

João Proença: Funding acquisition, Supervision, Formal analysis, Writing—review & editing.’

Ehsan Valiollahi: Formal analysis.

Vasco M. Barreto: Funding acquisition, Resources, Supervision, Formal analysis, Writing—original draft, review & editing.

## DATA AVAILABILITY

The vectors are available via Addgene (Plasmid #113096). The ICE web tool can be accessed at no cost through EditCo - CRISPR Performance Analysis. The Python scripts are available from the authors upon request.

