## Supplemental Data for "Endogenous intronic RNA tightly controls Cas9/CRISPR-mediated gene editing in human cells"

**SUPPLEMENTARY DATA**

Supplementary Table1. RNA molecules used in cgRNA experiments

| Oligo name | sequence |
|---|---|
| crRNA-intcgRNA 13 bp duplex | /AlTR1/rUrGrGrUrArArUrGrArUrGrGrCrUrUrCrArArCrArGrUrUrUrUrArGrArGrCrUrArUrGrCrUrCrArArUrCrUrGrGrGrUrArCrUrGrCrUrGrUrUrGrArArGrCrCrArUrCrGrArGrCrCrUrArGrCrCrUrCrArUrC/AlTR2/ |
| crRNA-intcgRNA 12 bp duplex | /AlTR1/rUrGrGrUrArArUrGrArUrGrGrCrUrUrCrArArCrArGrUrUrUrUrArGrArGrCrUrArUrGrCrUrCrArArUrCrUrGrGrGrUrArCrUrGrCrUrGrUrUrGrArArGrCrCrArUrGrArGrCrCrUrArGrCrCrUrCrArUrC/AlTR2/ |
| crRNA-intcgRNA 11 bp duplex | /AlTR1/rUrGrGrUrArArUrGrArUrGrGrCrUrUrCrArArCrArGrUrUrUrUrArGrArGrCrUrArUrGrCrUrCrArArUrCrUrGrGrGrUrArCrUrGrCrUrGrUrUrGrArArGrCrCrArGrArGrCrCrUrArGrCrCrUrCrArUrC/AlTR2/ |
| crRNA-native gRNA | /AlTR1/rUrGrGrUrArArUrGrArUrGrGrCrUrUrCrArArCrArGrUrUrUrUrArGrArGrCrUrArUrGrCrU/AlTR/ |
| Trigger RNA (not full length intron) | /AlTR1/rGrArUrGrArGrGrCrUrArGrGrCrUrCrUrGrGrArUrArUrCrUrGrCrArGrUrArCrCrCrArGrArUrUrG/AlTR2/ |

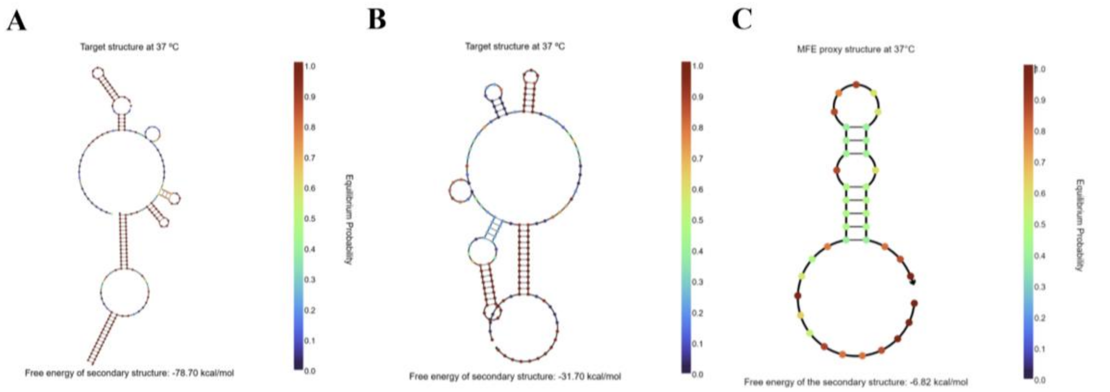

**Supplementary figure 1 . Predicted secondary structure and free energy of the intcgRNA states and trigger.** A) On-state secondary structure of the intcgRNA construct (proxy for the 13-bp blocking duplex version), predicted with NUPACK. B) Off-state secondary structure of the same intcgRNA construct. C) Predicted secondary structure of the intronic trigger RNA.

**Supplementary Table 2 . Transfection efficiency and post-sort purity in HPB-ALL cells.** $GFP^+$ percentages (transfection efficiency) were determined by flow cytometry 72 h after electroporation. Purity refers to the percentage of $GFP^+$ cells in the sorted population used for downstream Sanger sequencing analysis.

| HPB-ALL | | |
|---|---|---|
| Condition | Transfection Efficiency(%) | Purity(%) |
| Native gRNA + Cas9 | 31.7 | 98.3 |
| intcgRNA 11 + Cas9 | 51.2 | 96.0 |
| intcgRNA 12 + Cas9 | 37.0 | 96.8 |
| intcgRNA 13 + Cas9 | 29.3 | 95.9 |
| Native gRNA only | 15.3 | 98.1 |

**Supplementary Table 3 . Transfection efficiency and post-sort purity in HeLa cells.**GFP$^{+}$ percentages (transfection efficiency) were determined by flow cytometry 72 h after electroporation. Purity refers to the percentage of GFP$^{+}$ cells in the sorted population used for downstream Sanger sequencing analysis.

| Hela | | |
|---|---|---|
| Condition | Transfection Efficiency(%) | Purity(%) |
| Native gRNA + Cas9 | 42.5 | 98.8 |
| intcgRNA 11 + Cas9 | 41.6 | 98.5 |
| intcgRNA 12 + Cas9 | 43.1 | 97.6 |
| intcgRNA 13 + Cas9 | 47.6 | 98.3 |
| Native gRNA only | 41.2 | 97.8 |

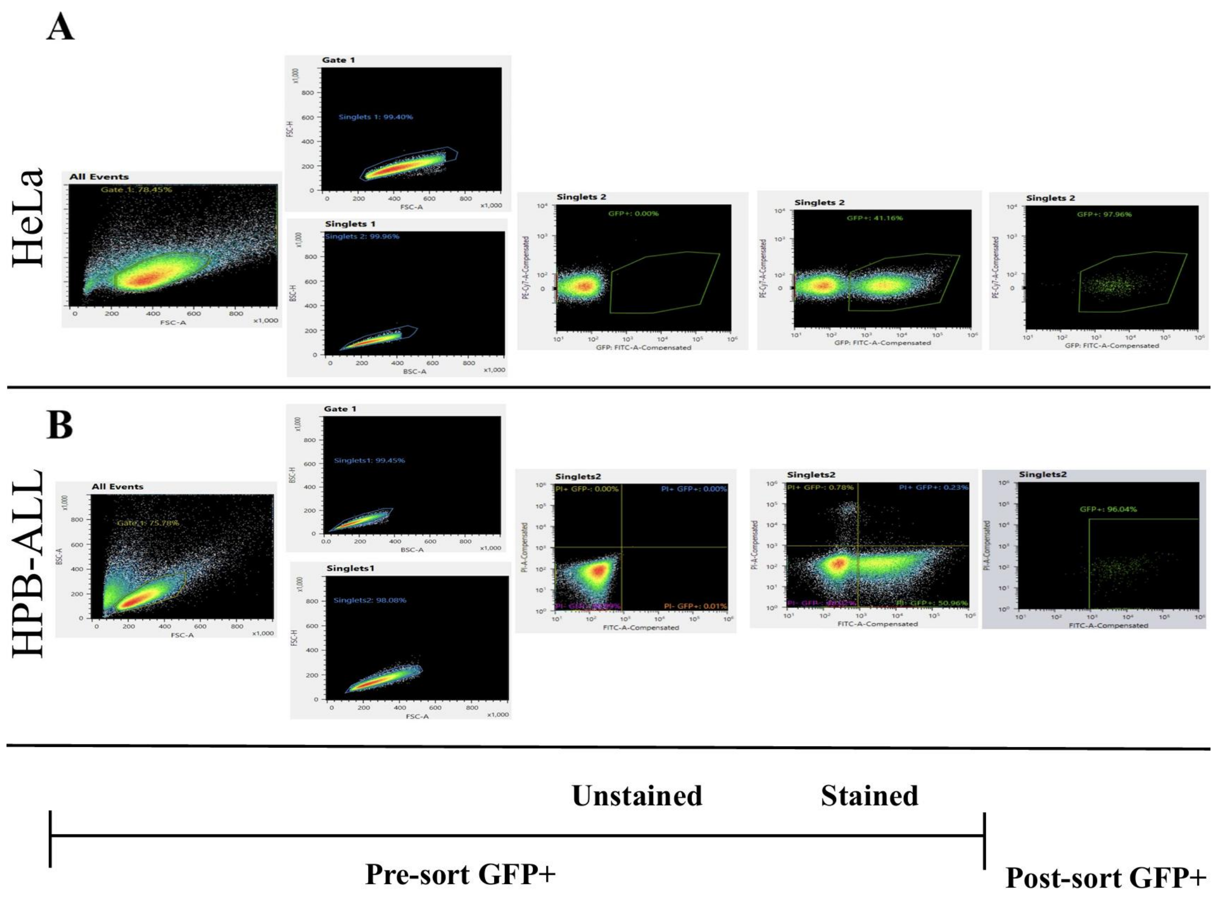


**Supplementary Figure 2 . Flow cytometry gating strategy and sorting purity for HeLa and HPB-ALL cells.** Representative flow cytometry plots showing the gating hierarchy used to identify and sort $GFP^+$ transfected cells 72 hours after electroporation with Cas9 RNP complexes and a GFP-encoding plasmid. (A) HeLa cells. (B) HPB-ALL cells. From left to right: forward/side scatter gating to exclude debris and dead cells (All Events, Gate 1), singlet discrimination ( Singlets 1, Singlets 2), followed by GFP fluorescence analysis on unstained controls, stained pre-sort populations, and post-sort $GFP^+$ populations. Post-sort purities exceeded 95% for all conditions (see Supplementary Tables 2 and 3). GFP was detected with a 488 nm laser and a 525/50 band-pass filter; PI viability staining (HPB-ALL) was detected using a 488 nm laser and a 600/60 band-pass filter.

A.

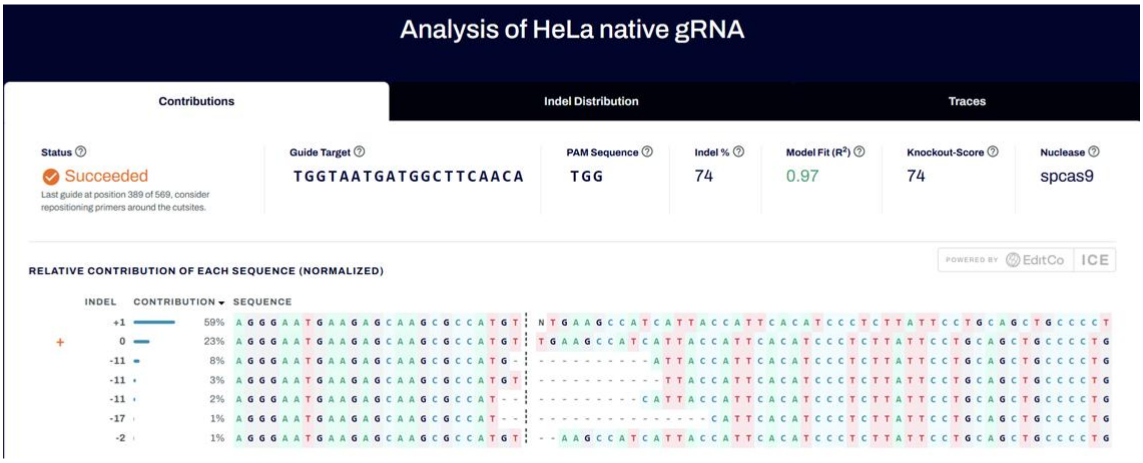

Analysis of HeLa native gRNA
Contributions
Indel Distribution
Traces
Status
Succeeded
Last guide at position 389 of 569, consider repositioning primers around the cutsites.
Guide Target
TGGTAATGATGGCTTCAACA
PAM Sequence
TGG
Indel %
74
Model Fit (R²)
0.97
Knockout-Score
74
Nuclease
spcas9
POWERED BY EditCo ICE
RELATIVE CONTRIBUTION OF EACH SEQUENCE (NORMALIZED)
INDEL
CONTRIBUTION
SEQUENCE
+1 59%
0 23%
-11 8%
-11 3%
-11 2%
-17 1%
-2 1%


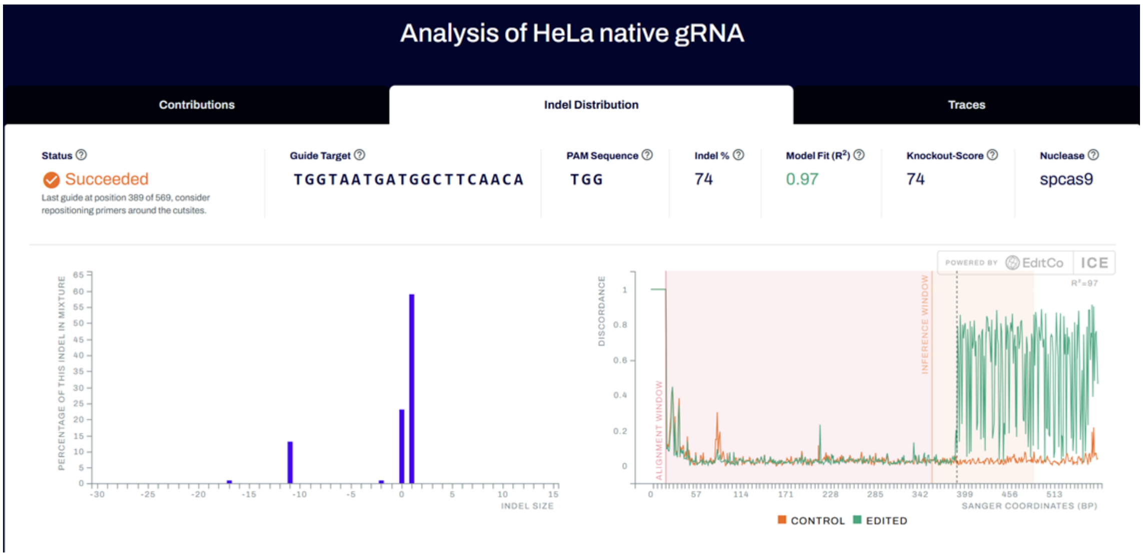

Analysis of HeLa native gRNA
Contributions
Indel Distribution
Traces
Status
Succeeded
Last guide at position 389 of 569, consider repositioning primers around the cutsites.
Guide Target
TGGTAATGATGGCTTCAACA
PAM Sequence
TGG
Indel %
74
Model Fit (R²)
0.97
Knockout-Score
74
Nuclease
spcas9
POWERED BY EditCo ICE
PERCENTAGE OF THIS INDEL IN MIXTURE
INDEL SIZE
DISCORDANCE
ALIGNMENT WINDOW
INFERENCE WINDOW
R²=97
SANGER COORDINATES (BP)
CONTROL
EDITED


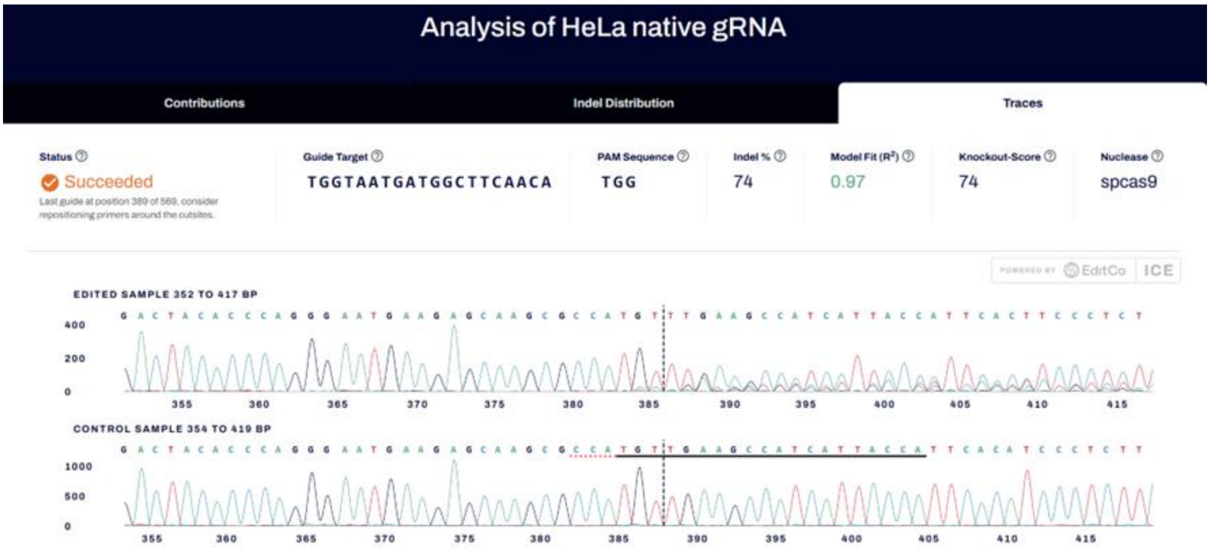

Analysis of HeLa native gRNA
Contributions
Indel Distribution
Traces
Status
Succeeded
Last guide at position 389 of 569, consider repositioning primers around the cutsites.
Guide Target
TGGTAATGATGGCTTCAACA
PAM Sequence
TGG
Indel %
74
Model Fit (R²)
0.97
Knockout-Score
74
Nuclease
spcas9
POWERED BY EditCo ICE
EDITED SAMPLE 352 TO 417 BP
CONTROL SAMPLE 354 TO 419 BP

B.

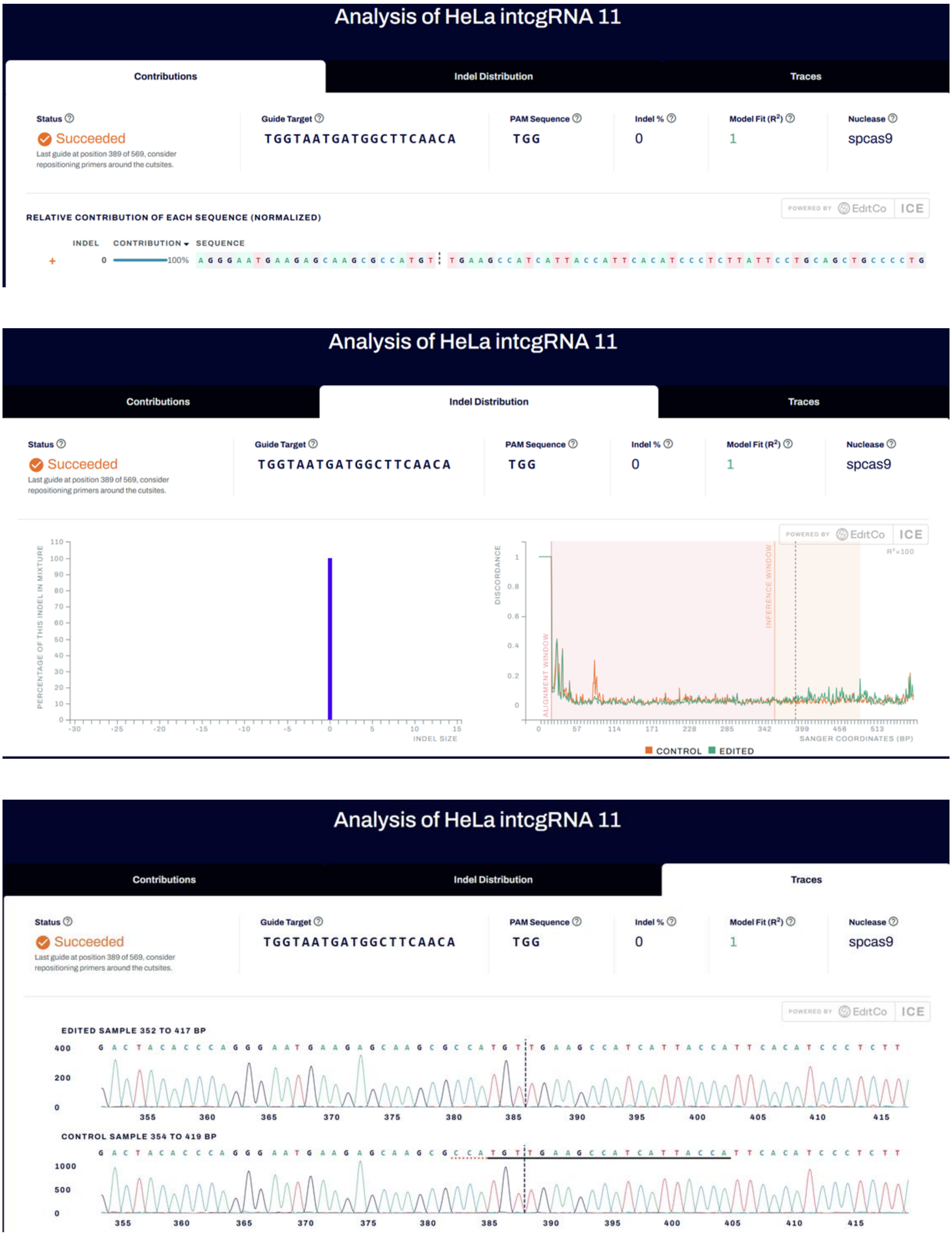

Analysis of HeLa intcgRNA 11
Contributions
Indel Distribution
Traces
Status
Succeeded
Last guide at position 389 of 569, consider repositioning primers around the cutsites.
Guide Target
TGGTAATGATGGCTTCAACA
PAM Sequence
TGG
Indel %
0
Model Fit (R²)
1
Nuclease
spcas9
POWERED BY EditCo ICE
RELATIVE CONTRIBUTION OF EACH SEQUENCE (NORMALIZED)
INDEL
CONTRIBUTION
SEQUENCE
0
100%
PERCENTAGE OF THIS INDEL IN MIXTURE
INDEL SIZE
DISCORDANCE
ALIGNMENT WINDOW
INFERENCE WINDOW
SANGER COORDINATES (BP)
R²=100
CONTROL
EDITED
EDITED SAMPLE 352 TO 417 BP
CONTROL SAMPLE 354 TO 419 BP

C.

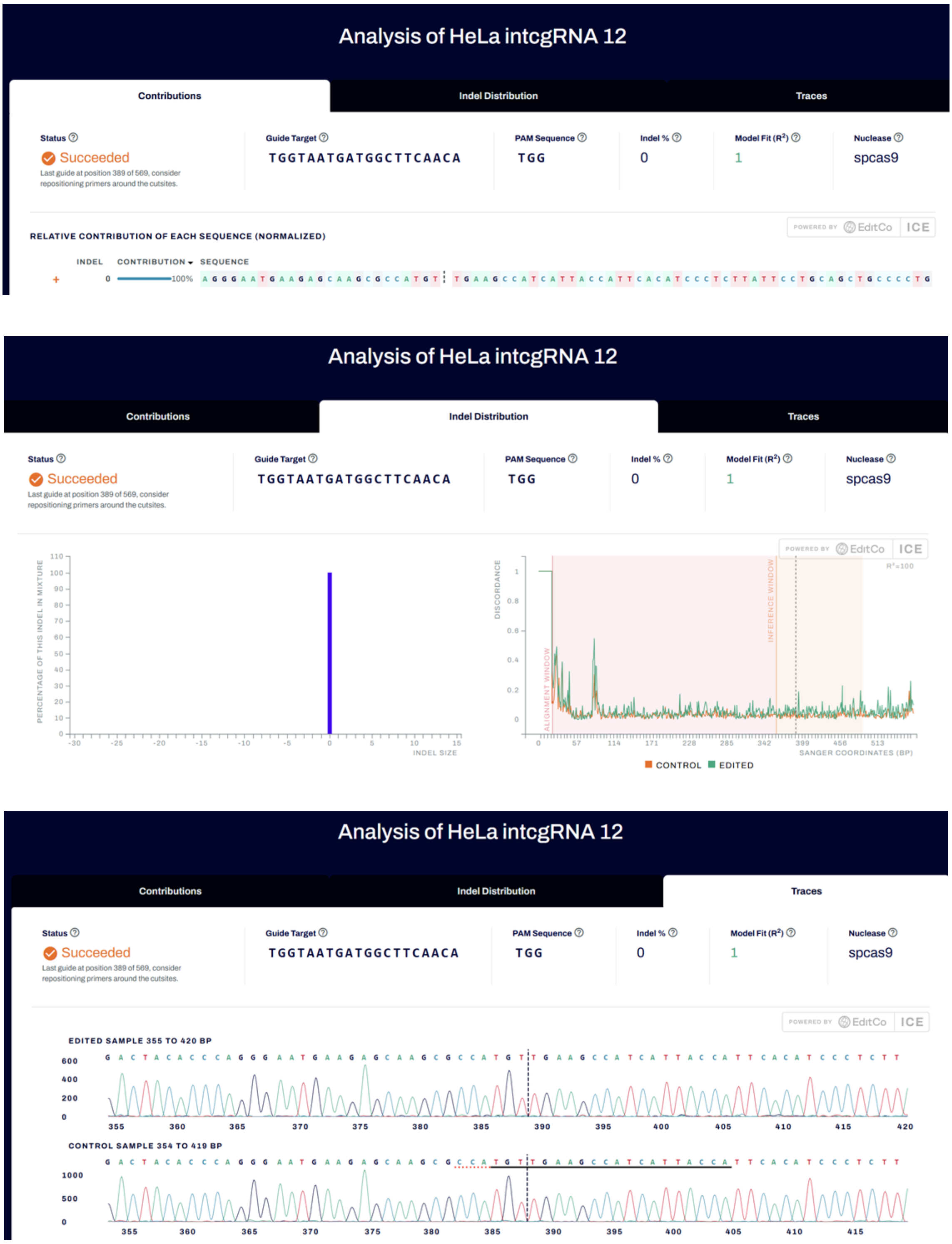
Analysis of HeLa intcgRNA 12
Contributions
Indel Distribution
Traces
Status
Succeeded
Last guide at position 389 of 569, consider repositioning primers around the cutsites.
Guide Target
TGGTAATGATGGCTTCAACA
PAM Sequence
TGG
Indel %
0
Model Fit (R²)
1
Nuclease
spcas9
POWERED BY EditCo ICE
RELATIVE CONTRIBUTION OF EACH SEQUENCE (NORMALIZED)
INDEL
CONTRIBUTION
SEQUENCE
0
100%
AGGGAATGAAGAGCAAGCGCCATGT | TGAAGCCATCATTACCATTCACATCCCTCTTATTCCTGCAGCTGCCCCTG
PERCENTAGE OF THIS INDEL IN MIXTURE
INDEL SIZE
DISCORDANCE
ALIGNMENT WINDOW
INFERENCE WINDOW
SANGER COORDINATES (BP)
R²=100
CONTROL
EDITED
EDITED SAMPLE 355 TO 420 BP
CONTROL SAMPLE 354 TO 419 BP

D.

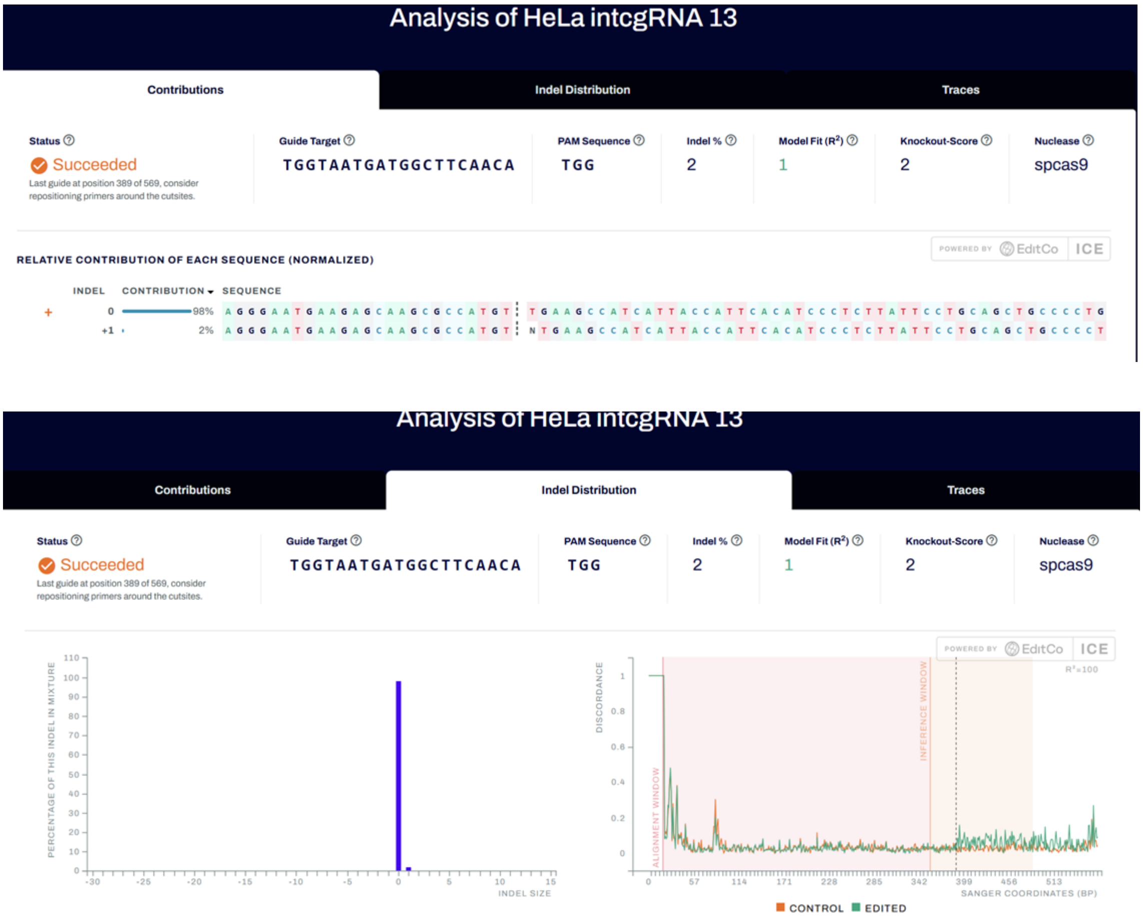

Analysis of HeLa intcgRNA 13
Contributions
Indel Distribution
Traces
Status
Succeeded
Last guide at position 389 of 569, consider repositioning primers around the cutsites.
Guide Target
TGGTAATGATGGCTTCAACA
PAM Sequence
TGG
Indel %
2
Model Fit (R²)
1
Knockout-Score
2
Nuclease
spcas9
POWERED BY EditCo ICE
RELATIVE CONTRIBUTION OF EACH SEQUENCE (NORMALIZED)
INDEL
CONTRIBUTION
SEQUENCE
0
98%
+1
2%
Analysis of HeLa intcgRNA 13
PERCENTAGE OF THIS INDEL IN MIXTURE
INDEL SIZE
DISCORDANCE
ALIGNMENT WINDOW
INFERENCE WINDOW
SANGER COORDINATES (BP)
CONTROL
EDITED

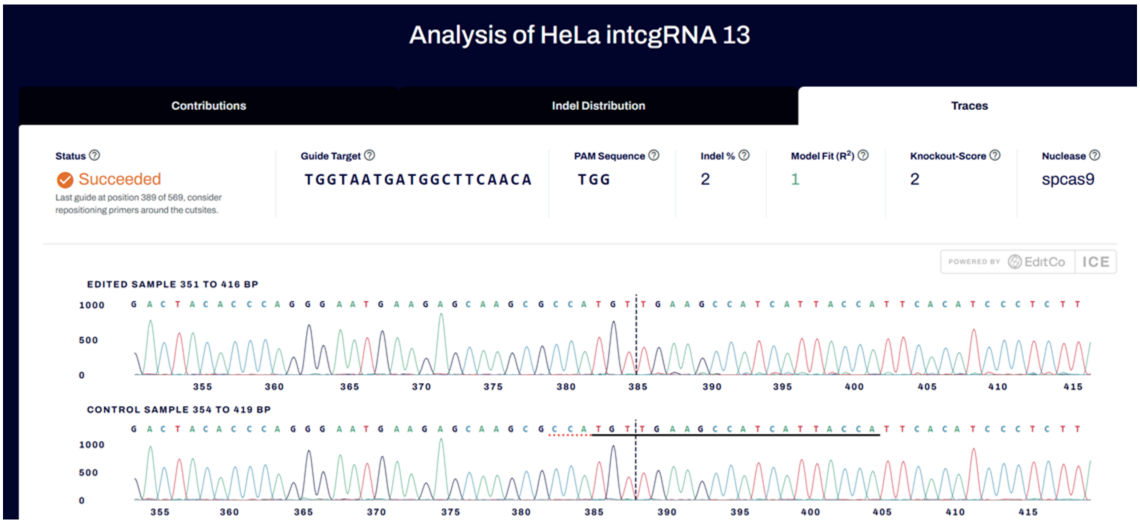

Analysis of HeLa intcgRNA 13
Contributions
Indel Distribution
Traces
Status
Succeeded
Last guide at position 389 of 569, consider repositioning primers around the cutsites.
Guide Target
TGGTAATGATGGCTTCAACA
PAM Sequence
TGG
Indel %
2
Model Fit (R²)
1
Knockout-Score
2
Nuclease
spcas9
POWERED BY EditCo ICE
EDITED SAMPLE 351 TO 416 BP
CONTROL SAMPLE 354 TO 419 BP


E.

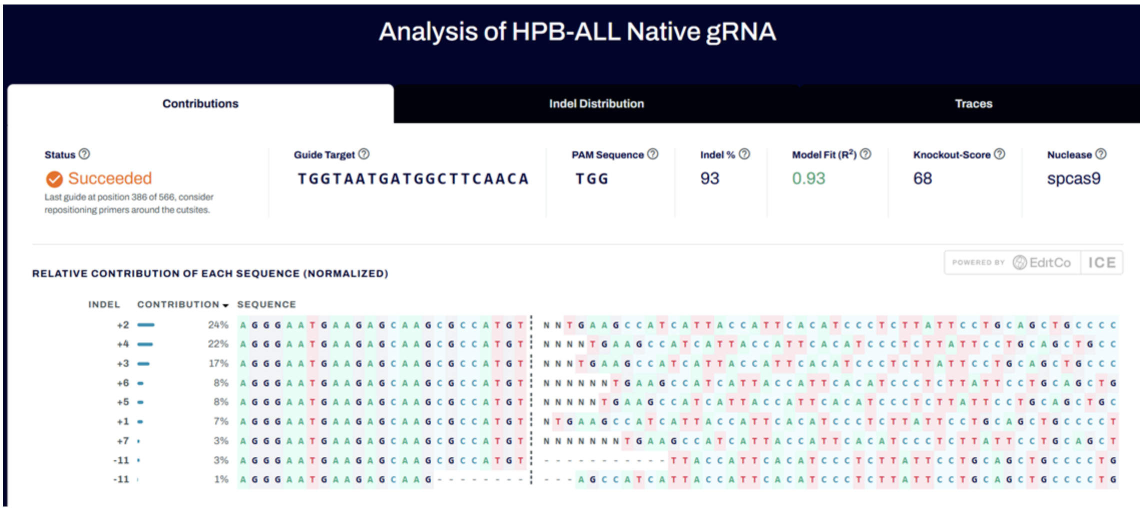

Analysis of HPB-ALL Native gRNA
Contributions
Indel Distribution
Traces
Status
Succeeded
Last guide at position 386 of 566, consider repositioning primers around the cutsites.
Guide Target
TGGTAATGATGGCTTCAACA
PAM Sequence
TGG
Indel %
93
Model Fit (R²)
0.93
Knockout-Score
68
Nuclease
spcas9
POWERED BY EditCo ICE
RELATIVE CONTRIBUTION OF EACH SEQUENCE (NORMALIZED)
INDEL
CONTRIBUTION
SEQUENCE
+2 24%
+4 22%
+3 17%
+6 8%
+5 8%
+1 7%
+7 3%
-11 3%
-11 1%

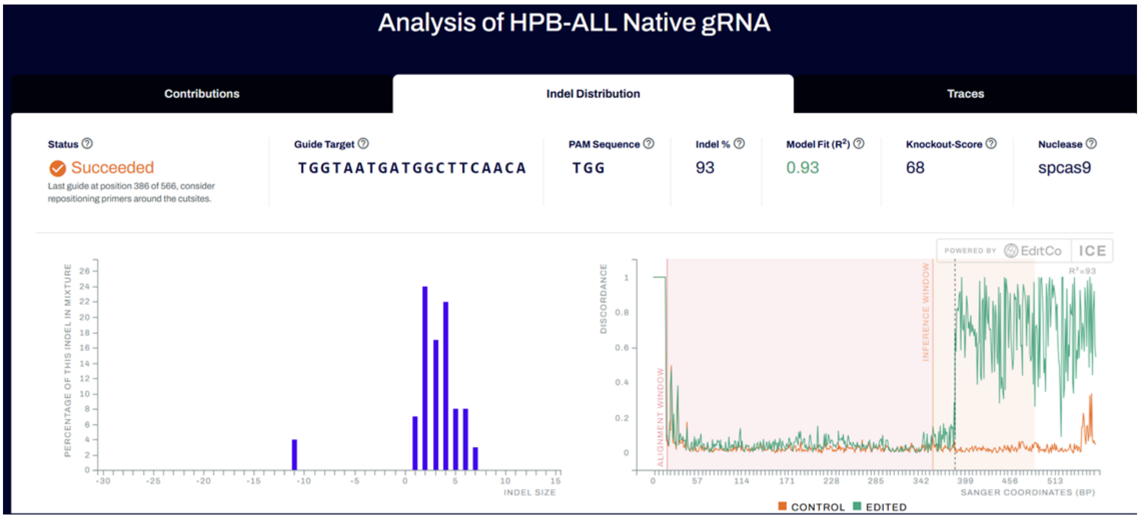
Analysis of HPB-ALL Native gRNA
Contributions
Indel Distribution
Traces
Status
Succeeded
Last guide at position 386 of 566, consider repositioning primers around the cutsites.
Guide Target
TGGTAATGATGGCTTCAACA
PAM Sequence
TGG
Indel %
93
Model Fit (R²)
0.93
Knockout-Score
68
Nuclease
spcas9
PERCENTAGE OF THIS INDEL IN MIXTURE
INDEL SIZE
DISCORDANCE
ALIGNMENT WINDOW
INFERENCE WINDOW
SANGER COORDINATES (BP)
CONTROL
EDITED
POWERED BY EditCo ICE

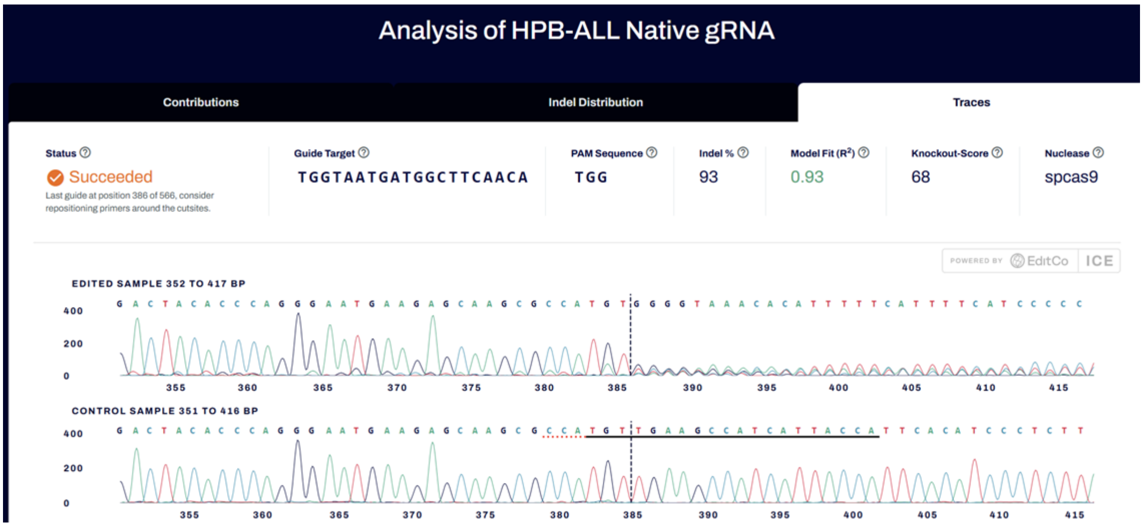
Analysis of HPB-ALL Native gRNA
Contributions
Indel Distribution
Traces
Status
Succeeded
Last guide at position 386 of 566, consider repositioning primers around the cutsites.
Guide Target
TGGTAATGATGGCTTCAACA
PAM Sequence
TGG
Indel %
93
Model Fit (R²)
0.93
Knockout-Score
68
Nuclease
spcas9
POWERED BY EditCo ICE
EDITED SAMPLE 352 TO 417 BP
CONTROL SAMPLE 351 TO 416 BP

F.

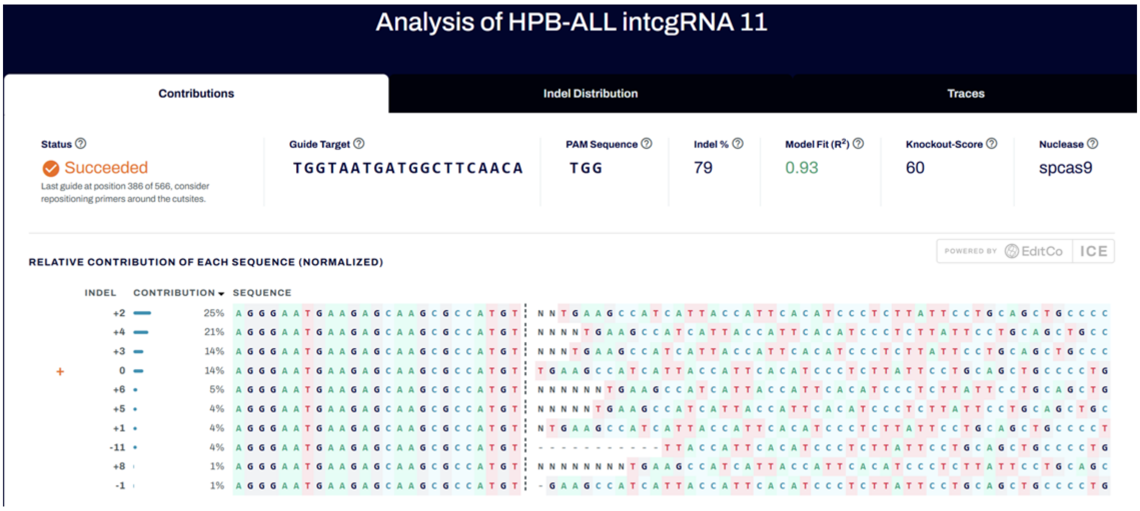
Analysis of HPB-ALL intcgRNA 11
Contributions
Indel Distribution
Traces
Status
Succeeded
Last guide at position 386 of 566, consider repositioning primers around the cutsites.
Guide Target
TGGTAATGATGGCTTCAACA
PAM Sequence
TGG
Indel %
79
Model Fit (R²)
0.93
Knockout-Score
60
Nuclease
spcas9
RELATIVE CONTRIBUTION OF EACH SEQUENCE (NORMALIZED)
INDEL
CONTRIBUTION
SEQUENCE
+2 25%
+4 21%
+3 14%
0 14%
+6 5%
+5 4%
+1 4%
-11 4%
+8 1%
-1 1%

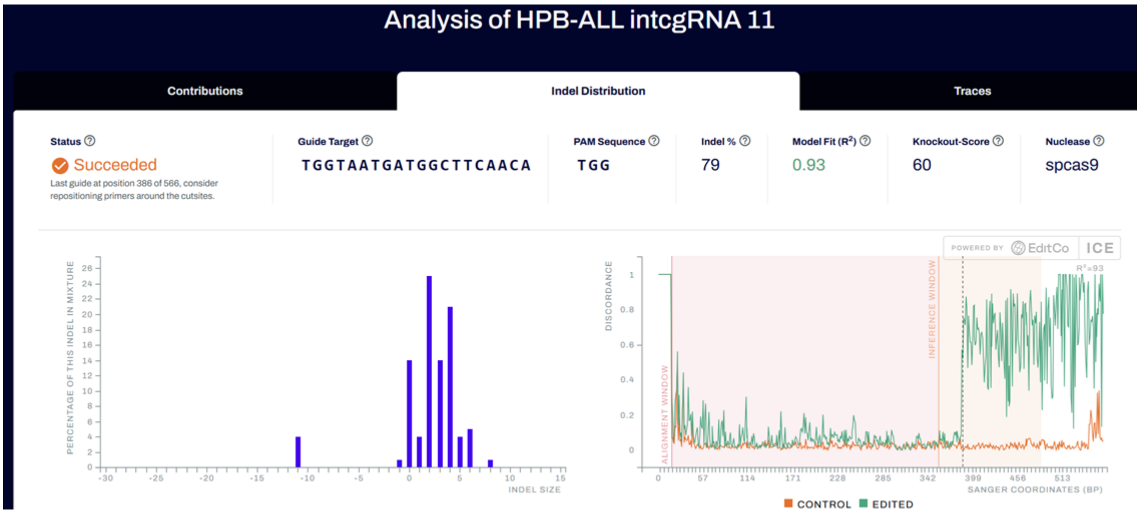
Analysis of HPB-ALL intcgRNA 11
Contributions
Indel Distribution
Traces
Status
Succeeded
Last guide at position 386 of 566, consider repositioning primers around the cutsites.
Guide Target
TGGTAATGATGGCTTCAACA
PAM Sequence
TGG
Indel %
79
Model Fit (R²)
0.93
Knockout-Score
60
Nuclease
spcas9
PERCENTAGE OF THIS INDEL IN MIXTURE
INDEL SIZE
DISCORDANCE
ALIGNMENT WINDOW
INFERENCE WINDOW
SANGER COORDINATES (BP)
CONTROL
EDITED

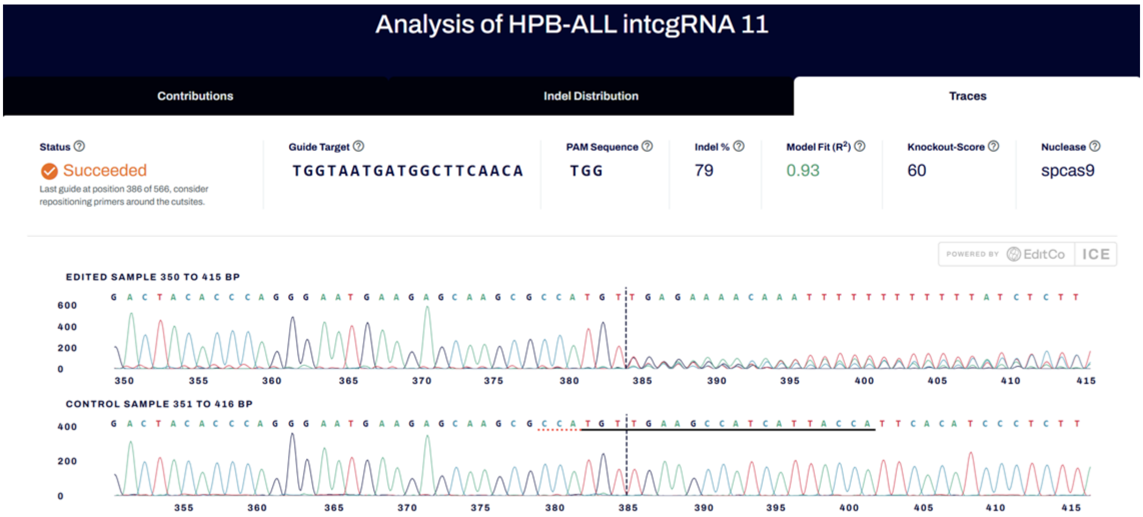
Analysis of HPB-ALL intcgRNA 11
Contributions
Indel Distribution
Traces
Status
Succeeded
Last guide at position 386 of 566, consider repositioning primers around the cutsites.
Guide Target
TGGTAATGATGGCTTCAACA
PAM Sequence
TGG
Indel %
79
Model Fit (R²)
0.93
Knockout-Score
60
Nuclease
spcas9
EDITED SAMPLE 350 TO 415 BP
CONTROL SAMPLE 351 TO 416 BP

G.

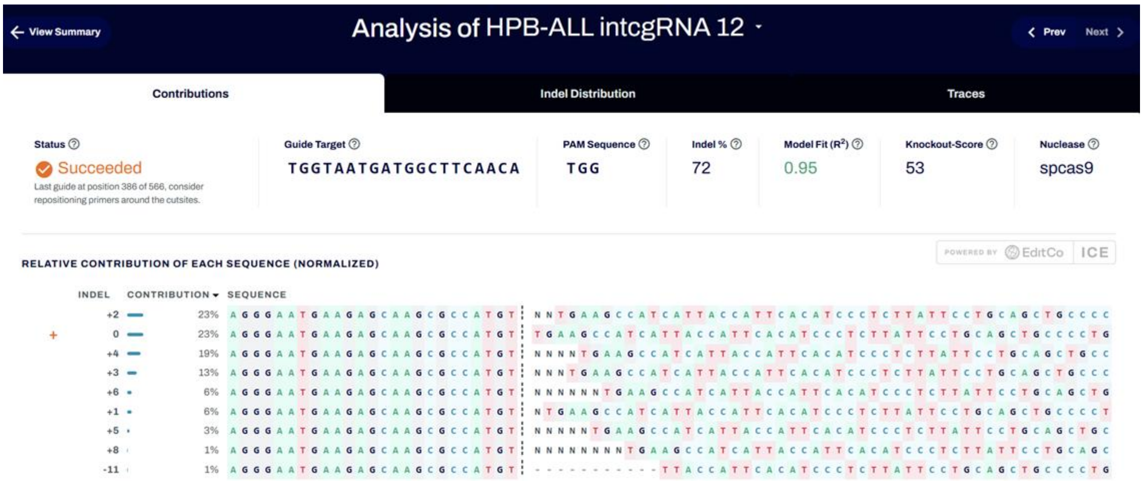
View Summary
Analysis of HPB-ALL intcgRNA 12
Prev
Next
Contributions
Indel Distribution
Traces
Status
Succeeded
Last guide at position 386 of 566, consider repositioning primers around the cutsites.
Guide Target
TGGTAATGATGGCTTCAACA
PAM Sequence
TGG
Indel %
72
Model Fit (R²)
0.95
Knockout-Score
53
Nuclease
spcas9
POWERED BY EditCo ICE
RELATIVE CONTRIBUTION OF EACH SEQUENCE (NORMALIZED)
INDEL
CONTRIBUTION
SEQUENCE
+2 23%
0 23%
+4 19%
+3 13%
+6 6%
+1 6%
+5 3%
+8 1%
-11 1%

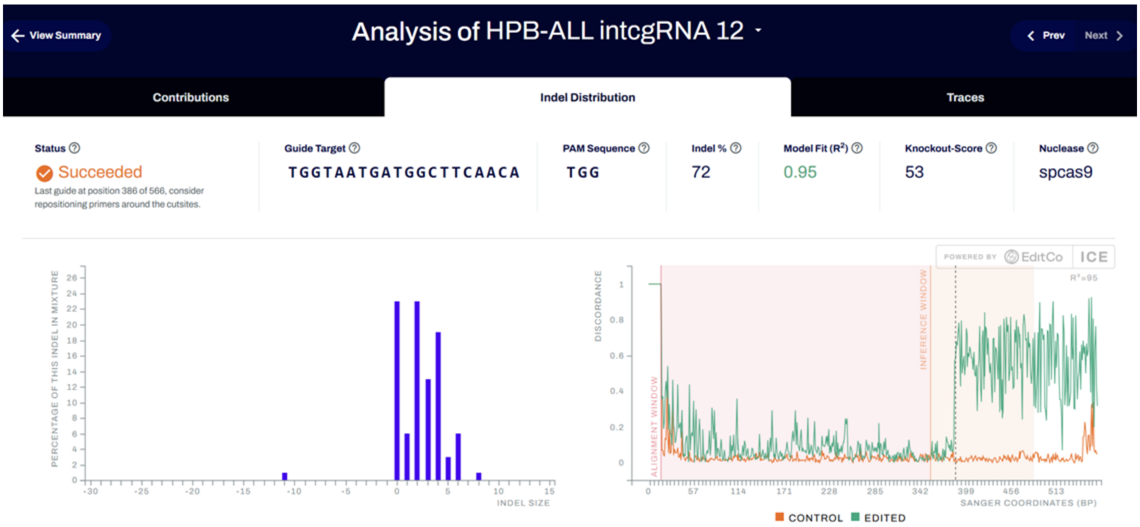
View Summary
Analysis of HPB-ALL intcgRNA 12
Prev
Next
Contributions
Indel Distribution
Traces
Status
Succeeded
Last guide at position 386 of 566, consider repositioning primers around the cutsites.
Guide Target
TGGTAATGATGGCTTCAACA
PAM Sequence
TGG
Indel %
72
Model Fit (R²)
0.95
Knockout-Score
53
Nuclease
spcas9
POWERED BY EditCo ICE
PERCENTAGE OF THIS INDEL IN MIXTURE
INDEL SIZE
DISCORDANCE
ALIGNMENT WINDOW
INFERENCE WINDOW
R²=95
SANGER COORDINATES (BP)
CONTROL
EDITED

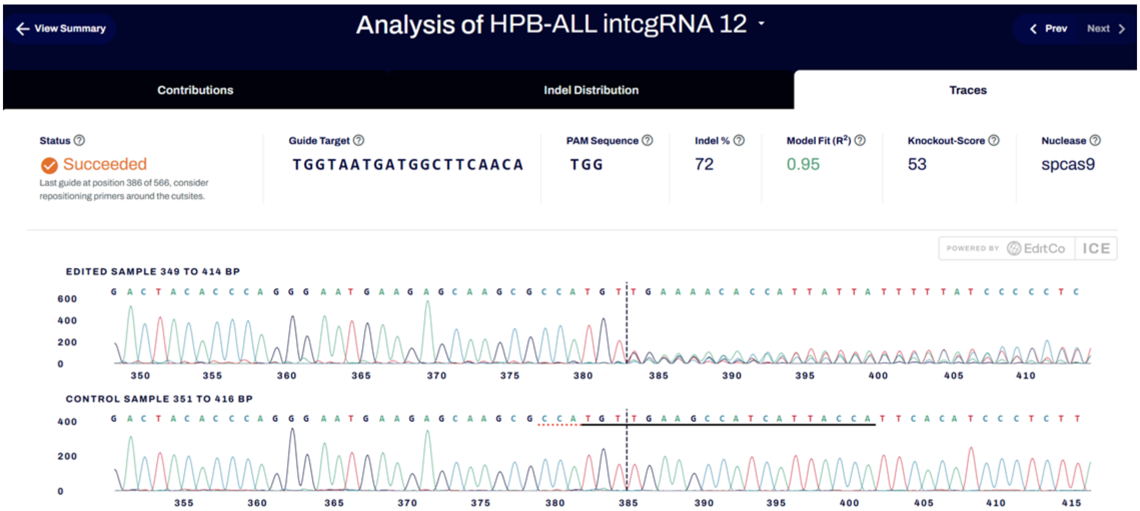
View Summary
Analysis of HPB-ALL intcgRNA 12
Prev
Next
Contributions
Indel Distribution
Traces
Status
Succeeded
Last guide at position 386 of 566, consider repositioning primers around the cutsites.
Guide Target
TGGTAATGATGGCTTCAACA
PAM Sequence
TGG
Indel %
72
Model Fit (R²)
0.95
Knockout-Score
53
Nuclease
spcas9
POWERED BY EditCo ICE
EDITED SAMPLE 349 TO 414 BP
CONTROL SAMPLE 351 TO 416 BP

H.

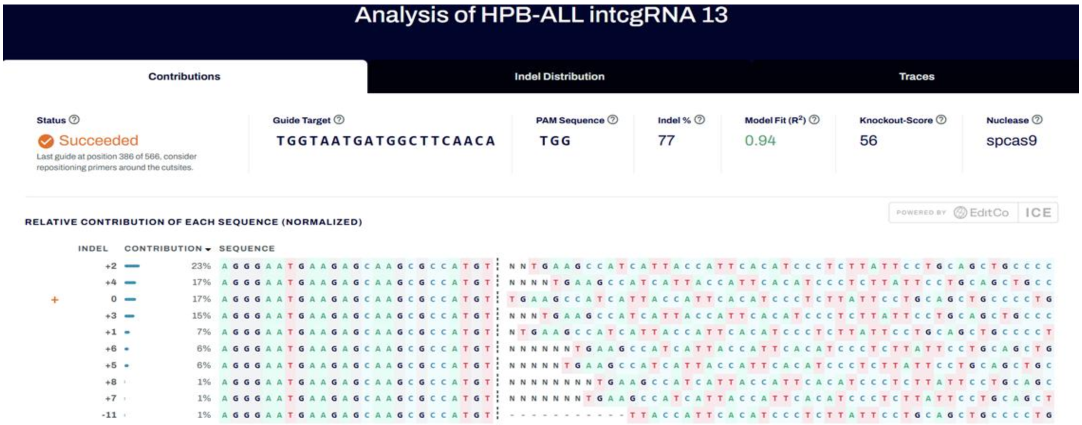


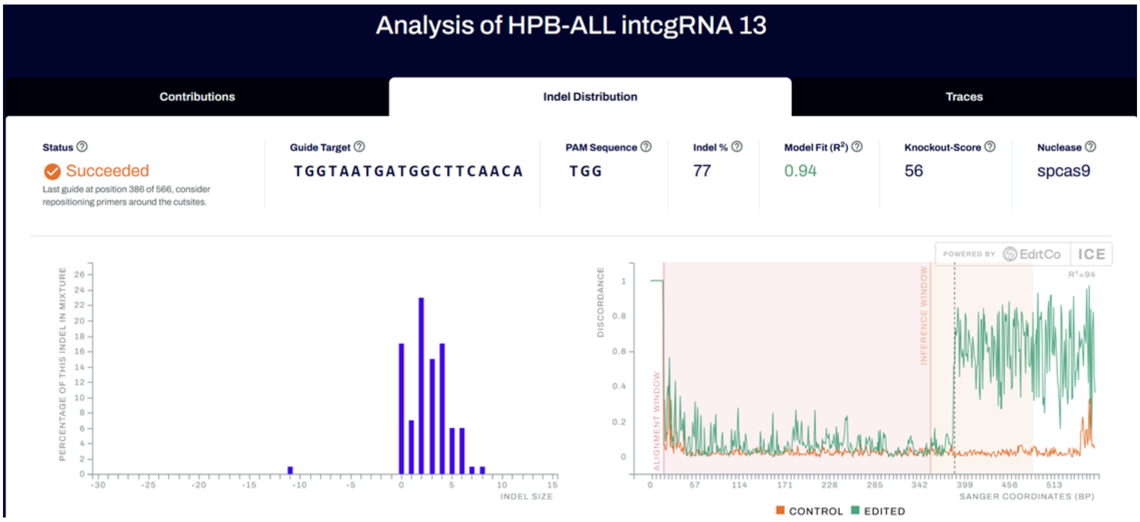


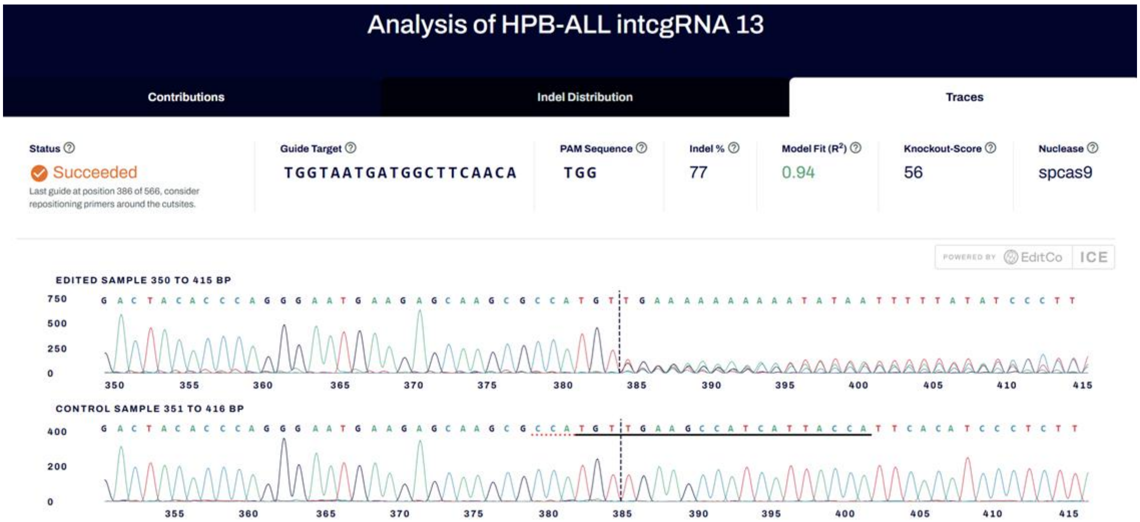


**Supplementary Figure 3. Representative Synthego ICE analysis outputs for all experimental conditions.** ICE trace decomposition, indel distribution, and quality metrics are shown for each RNP condition in HeLa (A–D) and HPB-ALL (E–H) cells. (A, E) Native crRNA; (B, F) intcgRNA with 11-bp duplex; (C, G) intcgRNA with 12-bp duplex; (D, H) intcgRNA with 13-bp duplex."
